# *Solanum galapagense*-derived purple tomato fruit color is conferred by novel alleles of the *Anthocyanin fruit* and *atroviolacium loci*

**DOI:** 10.1101/2021.11.09.467926

**Authors:** Sean Fenstemaker, Leah Sim, Jessica Cooperstone, David Francis

## Abstract

One hypothesis for the origin of endemic species of tomato on the Galápagos islands postulates a hybridization of *Solanum pimpinellifolium* and *S. habrochaites. S. galapagense* accession LA1141 has purple fruit pigmentation which has previously been described in green-fruited wild tomatoes such as *S. habrochaites*. Characterization of LA1141 derived purple pigmentation provides a test of the hybridization hypothesis. Purple pigmentation was recovered in progenies derived from LA1141 and the anthocyanins malvidin 3(coumaroyl)rutinoside-5-glucoside, petunidin 3-(coumaroyl) rutinoside-5-glucoside, and petunidin 3-(caffeoyl)rutinoside-5-glucoside were abundant. Fruit color was evaluated in an introgression population and three quantitative trait loci (QTLs) were mapped and validated in subsequent populations. The loci *atroviolacium* on chromosome 7, *Anthocyanin fruit* on chromosome 10, and *uniform ripening* also on chromosome 10, underly these QTLs. Sequence analysis suggested that the LA1141 alleles of *Aft* and *atv* are unique relative to those previously described from *S. chilense* accession LA0458 and *S. cheesmaniae* accession LA0434, respectively. Phylogenetic analysis of the LA1141 *Aft* genomic sequence did not support a green-fruited origin and the locus clustered with members of the red-fruited tomato clade. The LA1141 allele of *Aft* is not the result of an ancient introgression and underlies a gain of anthocyanin pigmentation in the red-fruited clade.

**Highlight:** *Anthocyanin fruit* and *atroviolacium* confer purple pigmentation in *Solanum galapagense* LA1141 confirming a mechanism described for green-fruited tomatoes. LA1141 alleles cluster with red-fruited homologs suggesting an independent gain of pigmentation.

## Introduction

Rick (1961) hypothesized that species of tomato endemic to the Galápagos, *L. cheesmanii f. minor* now classified as *Solnaum galapagense*, might have resulted from the hybridization of *S. pimpinellifolium* and *S. habrochaites* progenitors. This hypothesis was based on three unique traits found in both *S. habrochaites* and *S. galapagense*, including alleles of *B* capable of conferring high β-carotene (Lincoln and Porter, 1950; Tomes et al., 1954). *S. galapagense* also possesses accrescent calyx and pubescence reminiscent of *S. habrochaites* (Rick, 1961). *S. galapagense* accession LA1141 has purple pigmentation in immature fruit, similar to species in the green-fruited tomato clade including *S. habrochaites*. The presence of this fourth trait common to *S. galapagense* and *S. habrochaites* suggested that characterizing the chemical and genetic basis of purple fruit derived from LA1141 could provide a test of Rick’s 1961 hypothesis.

The endemic Galápagos tomatoes possess morphological and physiological traits that distinguish them from other wild species. These traits include orange fruit color at maturity, yellow-green foliage, tiny seed size, seed dormancy, and affinity for dry conditions (Rick, 1961; Darwin et al., 2003). These species can hybridize easily with cultivated tomato, making them useful donors of novel alleles (Rick, 1961). There are several genes from Galápagos tomatoes that have been used in breeding contemporary varieties. An allele of *uniform ripening* (*u*) from *S. cheesmaniae* accession LA0428 is responsible for uniform distribution of light green pigmentation in immature fruits (Rick, 1967). Alleles confering *jointless* (*j*^*2*^) pedicel (Rick, 1956) and *arthritic articulation* (*j*^*2in*^) (Joubert, 1961) have enabled mechanical harvest. *S. cheesmaniae* accession LA0422 has a recessive allele, *anthocyanin gainer* (*ag*^*2*^), which results in fruit and foliage lacking anthocyanin at early developmental stages (Rick, 1967; De Jong et al., 2004). Alleles of the *Beta* (*B*) locus found in all *S. galapagense* and *S. cheesmaniae* accessions confer high β-carotene and the characteristic orange fruit (Orchard et al., 2021). Alleles of *B* from LA0317 and LA0166 have been introgressed into cultivated tomatoes (Stommel, 2001). Anthocyanin-mediated purple pigmentation in both the fruit and foliage was described in *S. cheesmaniae* accession LA0434, the donor of the *atroviolacea* (*atv*) locus (Rick 1956; Rick, 1961; Rick, 1967). Additionally, an accession of *S. cheesmaniae* (LA0428) was described as having immature fruits with a purple color that resemble *S. peruvianum* (Rick, 1967). Identification and analysis of loci that confer purple fruit color may shed light on broader questions about the evolutionary history of the Galápagos tomatoes.

Water-soluble vacuolar pigments called anthocyanins cause purple fruit pigmentation in species of *Solanum* (Timberlake and Bridle, 1982; Mes et al., 2008; Chaves-Silva et al., 2018). The red-fruited tomato clade corresponds to the group *Lycopersicon* which generally lack anthocyanins in the fruit. The green-fruited clade is grouped into *Arcanum, Eriopersicon* and *Neolycopersicon* based whole genome sequence phylogeny (The 100 Tomato Genome Sequencing Consortium, et al., 2014). Purple pigmentation is a characteristic found throughout the green-fruited tomato clade. As an example, *S. habrochaites* accession LA1777 has pronounced anthocyanin spots in its fruit (Dal Cin et al., 2009). Additionally, *S. peruvianum* fruit are purple tinged with purple lines and blotches (Muller, 1940). The chemical basis of purple fruit derived from tomato species in the green-fruited clade is attributed to the anthocyanins petunidin and malvidin (Jones et al., 2003; Mathews et al., 2003, Ooe et al., 2016). Two loci are known to affect the regulation of anthocyanin accumulation in tomato fruit, one on chromosome 7 and a second on chromosome 10. A nonfunctional R3 MYB repressor on chromosome 7 underlies the *atv* locus (Cao et al., 2017). On chromosome 10, a functional R2R3 MYB-encoding activator gene underlies the *Anthocyanin fruit* (*Aft*) locus described in the donor parent *S. chilense* accession LA0458 (Georgiev, 1972 Jones et al., 2003; Mes et al., 2008). Additionally, the *aubergine* (*Abg*) locus from *S. lycopersicoides* accession LA2408 results in dark purple fruit (Rick et al., 1994). The *Abg* locus also maps to chromosome 10 and may be allelic to *Aft* (Rick et al., 1994). The synergistic interaction between a nonfunctional R3 MYB repressor *atv* and a functional R2R3 MYB activator at *Aft* elevates anthocyanin levels in tomato fruit and imparts purple color (Povero et al., 2011; Colanero et al., 2020 a Yan et al., 2020).

We conducted experiments aimed at describing the chemical and genetic basis of purple pigmentation in fruit derived from LA1141. Our results are consistent with the regulatory mechanism described for accessions from the green-fruited tomato clade. However, the LA1141 alleles of *Aft* and *atv* are distinct from those previously characterized. Phylogenetic analysis of *Aft* sequence does not support a green-fruited origin of the LA1141 locus which suggests that purple fruit pigmentation in this accession is the result of a convergent or parallel mechanism resulting from a loss of function that disrupts *atv* and a gain of function that restores *Aft*. These findings fail to support Rick’s 1961 hypothesis on the origin of the Galápagos tomatoes.

## Materials and methods

### Plant materials and growing conditions

An inbred backcross (IBC) population was initiated in 2014 for the simultaneous introduction and characterization of purple pigmented fruit. The IBC population was derived from an initial hybridization of *Solanum galapagense* S.C. Darwin and Peralta (formerly *Lycopersicon cheesmaniae* f. minor) (Hook. f) C.H.Mull.) accession LA1141 as the female parent and *Solanum lycopersicum* L. (formerly *Lycopersicon esculentum* Mill) OH8245 as the male parent. Accession LA1141 was acquired from the C.M. Rick Tomato Genetic Resources Center, University of California, Davis, CA, USA. The processing tomato variety OH8245 was described previously (Berry et al., 1991). A single LA1141 × OH8245 F_1_ plant was backcrossed as the female parent to OH8245. BC_1_ individuals were then separately backcrossed again with OH8245 as the pollen donor. BC_2_ plants were then self-pollinated with single seed descent in alternating greenhouse and field production cycles to create a BC_2_S_3_ IBC population composed of 160 inbred backcross lines (IBLs). During these studies, the IBC population was further inbred to BC_2_S_5_. The BC_2_S_3_ IBLs SG18-124 (Fig. 1C) and SG18-200 (Fig. 1B) were selected based on purple pigmentation in the fruit to generate populations for validation of quantitative trait loci (QTLs). The IBLs SG18-124 and SG18-200 were again crossed to OH8245, and the self-pollination of the resulting F_1_ plants created populations with F_2_ segregation for specific LA1141 chromosomal regions.

**Fig. 1.**
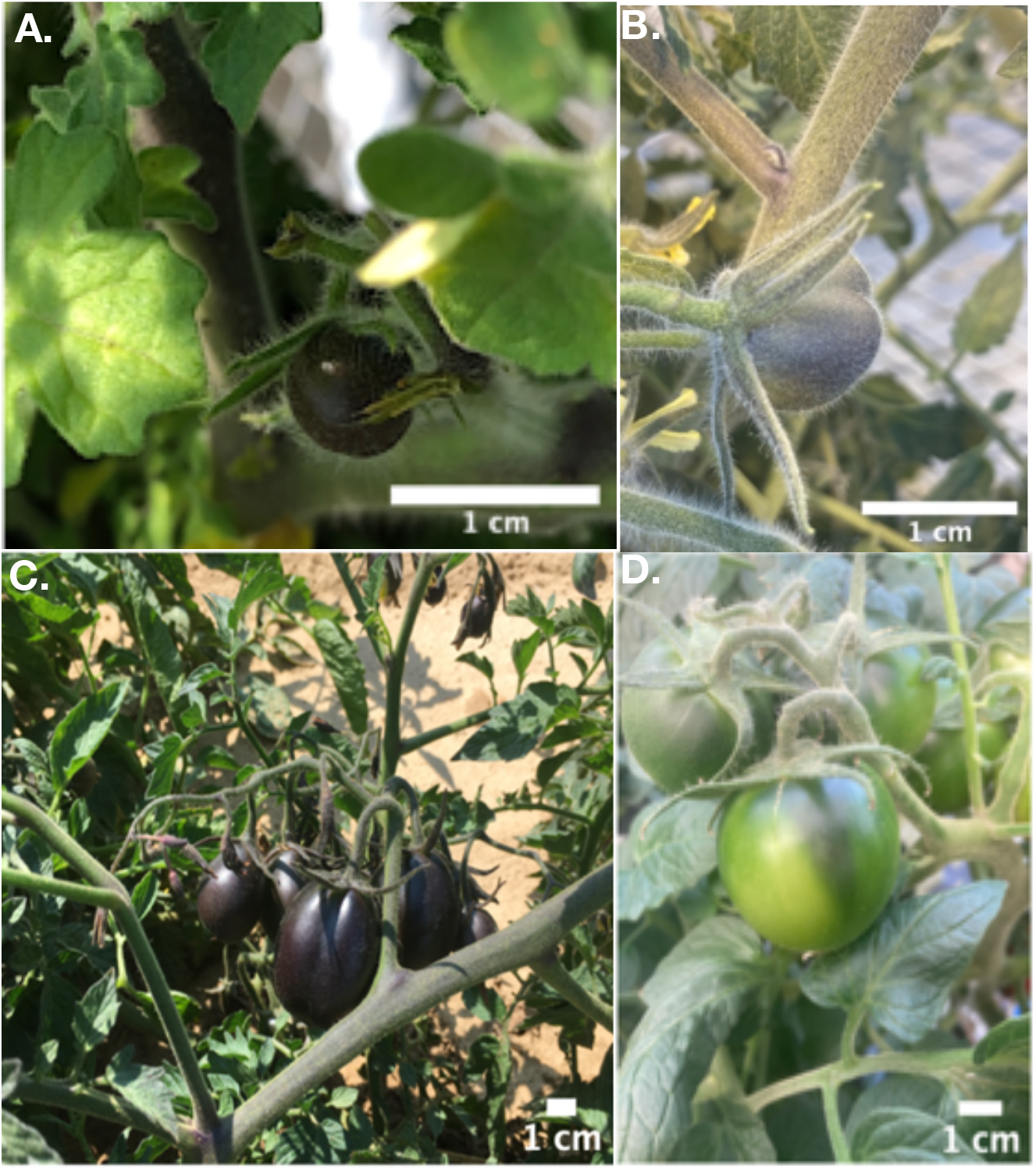
Heritable fruit pigmentation from *S. galapagense* accession LA1141. We determined a role for several candidate genes underlying the *Anthocyanin fruit* (*Aft*), *atroviolacea* (*atv*), and *uniform* ripening (*u*) loci derived from. Homozygous LA1141 *Aft* is designated as *Aft/Aft*, homozygous LA1141 *atv* is designated as *atv/atv*, and homozygous LA1141 *u* is designated as *U/U*. Notation follows previous publications Cao et al., 2017. **A**. LA1141 mature green fruit (*Aft*/*Aft*; *atv*/*atv*; *U*/*U*) **B**. Inbred backcross line (IBL) SG18-200 (*Aft/Aft; ATV/ATV; u/u*) **C**. IBL SG18-124 (*Aft/Aft; atv/atv*; U/U) **D**. IBL SG18-251 (*AFT/AFT*; *ATV/ATV*; *U/U*).

Seedling care for greenhouse and field trials followed the same protocol. The 160 BC_2_S_3_ IBLs and the SG18-124 and SG18-200 derived F_2_ progenies were sown in 288-cell trays with a cell volume of 13 ml. Greenhouse temperatures were set to 27 °C during the day and 25 °C at night with a 16-hour photoperiod. Photosynthetically active radiation (PAR) was supplied by natural sunlight, 1000-W metal-halide lamps (Multi-Vapor® GE Lighting, East Cleveland, OH, USA), and 1000-W high-pressure sodium lamps (Ultra Sun® Sunlight Supply, Vancouver, WA) with a target radiation of 250 W m^-2^ or approximately 113 µmol m^-2^ s^-1^ PAR. Fertilization was applied using a 20-20-20 fertilizer (20 percent N, 20 percent P_2_O_5_, and 20 percent K_2_O) (Jack’s professional All-Purpose Fertilizer, JR Peters INC., Allentown, PA, USA) delivered at a concentration of 1000 mg L^-1^ twice per week. Plants were irrigated once or twice per day as needed.

IBC and F_2_ progenies were evaluated in field trials as single plants. The BC_2_S_3_ IBC population was evaluated with 60 cm spacing and 164 plants, including controls. Progenies derived from SG18-124 and SG18-200 were transplanted for greenhouse and field evaluations of pigmentation in the fruit. Plants with three to five expanded leaves were transferred to 3.78 L containers (Hummert, EARTH City, MO) and spaced 30 cm apart on the greenhouse bench. There were 36 F_2_ plants evaluated in the greenhouse. The remaining SG18-124 and SG18-200 derived F_2_ progenies were evaluated in field trials with 60 cm spacing with a total of 145 plants harvested.

### Tomato Fruit Color Measurements

Three mature green fruit were randomly selected from each plot and measured at the midpoint between the shoulder and the blossom end of the fruit. Color was measured with a colorimeter (chromameter CR-300; Minolta Camera Co., Ltd., Ramsey, NJ, USA). Values of the red, green, yellow, and blue components of fruit were obtained using the “L*a*b*” CIELAB color space (Commission Internationale de l’Eclairage, 1978). The L* coordinate represented a measure of the darkness or lightness. Coordinates, a* and b*, are measured color along the axis of a color wheel with +a* in the red direction, and –a* in the green direction, +b* in the yellow direction, and –b* in the blue direction (Kabelka et al., 2004). Chroma and hue were derived from measurements of a* and b*. Chroma was calculated as (a*^2^ + b*^2^)^1/2^ and was used to measure of how bright or dull a color was. Hue was calculated using the equation (180/π) [cos^−1^ (a*/chroma)] for positive values of a*. For negative values of a*, we calculated hue using the equation 360-(180/π) [cos–1 (a*/chroma)] (Kabelka et al., 2004; Darrigues et al., 2008). The average values of hue, chroma, and L* were used as the response variable for our genetic studies.

### Anthocyanidin extraction, analysis, and identification

Tomato fruit samples at different stages of maturity from SG18-124 × OH8245 derived F_2_ plants were blended, and 3.5 g of juice was extracted with 4 ml 1% HCl in MeOH. The extracts were dried under nitrogen gas. Anthocyanins were separated using an C18 column (HSS T3, 2.1×100mm, 1.8um, Agilent Technologies) and a gradient of water (A) and acetonitrile (B), both with 5% formic acid. The gradient was as follows: isocratic with 0% B from 0-2 min, linear gradient to 30% B from 2-8 min, linear gradient to 100% from 8-12 min, hold at 100% B for 1 min, and return to initial conditions. Samples were run on an Agilent 1290 ultra-high-performance liquid chromatography (UHPLC) with photodiode array (PDA) detection, coupled to a high resolution 6545 quadrupole time-of-flight mass spectrometer (QTOF-MS) (Agilent, Santa Clara, CA, USA). The MS was run using electrospray ionization and operated in both positive and negative modes using reference masses for accurate mass determination.

### DNA isolation and genotyping

Genomic DNA was isolated from fresh, young leaf tissue from the 160 BC_2_S_3_ progenies, 96 of each F_2_ propulation, and parental lines using a modified CTAB method consistent with previous studies (Sim et al., 2015). Single-nucleotide polymorphisms (SNPs) between OH8245 and LA1141 were identified using a 384-marker panel (Bernal et al., 2020). Genotyping of the BC_2_S_3_ progenies was performed using the PlexSeq™ platform as a service (Agriplex Genomics, Cleveland, OH, USA) to detect specific SNPs through a pooled amplicon sequencing strategy.

### Marker development for candidate genes

Selected SNP markers and candidate genes were converted to polymerase chain reaction (PCR) based insertion/deletion (INDEL) markers for visualization on agarose gels. These markers, when appropriate, were added to the linkage maps described below. A summary containing forward and reverse primers, genome location, and expected polymorphism for these markers is available at https://doi.org/10.5281/zenodo.5650150 (Fenstemaker et al., 2021a). Candidate genes included MYB transcription factor sequences corresponding to *atv* [MF197509, NC_015447.3 (Cao et al., 2017)], *Aft* [EF433416; EF433417; MN433086; MN433087; FJ705319; NC_015447.3 (Mes et al., 2008; Sapir et al., 2008; Cao et al., 2017)], *GOLDEN2-LIKE (GLK2)* transcription factor sequences corresponding to *u* [JX163897; JQ316459; NC_015447.3 (Powell et al., 2012)], and *Lycopene β-cyclase (Cyc-B)* sequences corresponding to *B* [KP233161, (Orchard, 2014)]. These sequences were targeted as candidate genes based on initial quantitative trait locus (QTL) mapping and because of their previously known effects on tomato fruit color. The INDEL and cleaved amplified polymorphism sequences (CAPS) markers were developed using a sequence comparison approach between, *S. lycopersicum* variety Heinz 1706, *S. galapagense* accession LA1044, *Solanum cheesmaniae* (L.Riley) Fosberg, 1987 1) in [Fosberg FR (1987b)] (formerly *Lycopersicon cheesmaniae* L.Riley, 1925 in [Riley LAM (1925c)] accession LA0483, *S. cheesmaniae* accession LA1401, and *S. lycopersicum* variety Indigo Rose. Primers were designed using Primer3 (v.0.4.0) (Untergasser et al., 2012). These primers were used to genotype LA1141, OH8245, the BC_2_S_3_ IBC population, and the subsequent F_2_ progenies derived from IBL selections SG18-124 and SG18-200.

PCR was carried out with an initial incubation at 94 °C for 3 min, followed by 40 cycles of denaturation at 94 °C for 30 s, annealing at 52 °C for 30 s, and extension at 72 °C for 60 s. A final elongation step at 72 °C was carried out for 7 min after completing the cycles. The PCR products for markers detected as CAPS were digested with a restriction enzyme (Fenstemaker et al., 2021a) for two hours at 37 °C. The PCR products were separated on a 2.5% agarose gel.

### Linkage map construction

A genetic linkage map was constructed based on the IBC population. The R/qtl package version 1.47-9 was used in the R statistical software environment version 4.0.3 (Broman et al., 2003; R Core Team, 2020). We used the “read.cross” function from BC_s_F_t_ tools to read in data, with s = 2 and t = 3 (Shannon et al., 2013). Of the 384 SNPs in the marker panel, 157 were polymorphic in the IBC population, and no markers were removed. A summary of the 157 polymorphic SNPs is available at https://doi.org/10.5281/zenodo.5650152 (Fenstemaker et al., 2021b). The genetic map was constructed by using the “est.map” function in R/qtl. Markers were placed in the same linkage group if they had a logarithm of the odds (LOD) score greater than 1.8 and an estimated recombination fraction lower than 0.45. The Kosambi map function was used for map construction and to convert recombination frequency to genetic distance (Kosambi 1944). The marker order in each linkage group was estimated with the functions “orderMarkers” and “ripple”in R/qtl. Changes in chromosome length and loglikelihood were investigated, dropping one marker at a time with the function “droponemarker” in R/qtl. Marker order was compared to the physical position in the Tomato Genome version SL4.0 (Hosmani et al., 2019) using both linear (adjusted correlation coefficient R^2^) and rank regression (rho(ρ)) to assess linkage map quality.

### QTL Analysis in BC_2_S_3_ IBLs

Composite interval mapping (CIM) was used for QTL detection (Zeng, 1994) using the “cim” function in the R/qtl package (Broman et al., 2003). Analysis was performed using a 2 cM step, one marker selected as a cofactor, and a 40 cM window with cofactor and window selected due to limited recombination and expected skewed segregation in the BC_2_S_3_ population. Haley Knott regression (Haley and Knott, 1992) (hk) was chosen as the solution-generating algorithm. Significance thresholds were generated by using the permutation test (α = 0.05, n = 1000; Churchill and Doerge, 1994). The resampled LOD cutoffs used were LOD= 6.8 for hue, LOD = 4.5 for chroma, and LOD = 3.65 for L*. Genetic effects were evaluated as differences between phenotype averages expressed as regression coefficients using the “fitqtl” function with the argument “get.ests=TRUE” and “dropone=FALSE” in R/qtl. Additionally, percent variance explained was estimated by the “fitqtl” function with the argument “dropone=TRUE” in R/qtl.

### QTL validation

The IBLs SG18-124 and SG18-200 were chosen because of pigmentation in their fruit (Fig. 2B, C). Segregating SG18-124 × OH8245 F_2_ and SG18-200 × OH8256 F_2_ progenies were sown in the field and greenhouse, and fruit pigmentation was measured using the Minolta CR300 colorimeter as described above. Seedlings were also grown as previously described. Marker data were scored on 91 progenies derived from the SG18-124 × OH8245 F_2_ population and 90 from the SG18-200 × OH8245 F_2_ population. Genetic effects and allele substitutions were evaluated using linear model ANOVA as implemented by the “lm” function in the R core package (R Core Team, 2020). The linear model *Y* = *μ x* + *M*+ *E*: where *Y* was the color trait value, *μ x* was equal to the population mean, *M* was the effect of each marker allele, and *E* was the associated error, equivalent to genotype (marker). We compared the marker-locus genotypic classes of homozygous LA1141 *Aft (Aft*/*Aft*) and homozygous LA1141 *atv* (*atv*/*atv*), homozygous OH8245 *Aft* (*AFT/AFT)* and homozygous OH8245 *atv* (*ATV/ATV*), and all possible marker-locus class combinations. For consistency, the marker-locus genotypic class notation followed previous publications (Cao et al., 2017). The markers Ant1_1 (*Aft*), An2-like_exon2_intron2 (*Aft*), atv_ex4 (*atv*), u_gal_3 (*u*), and BetaRSA (*B*) were tested. The Marker evaluations were conducted in both F_2_ populations independently. We used F-tests as previously described to determine if hue, chroma, and L* were associated with significant differences in marker-locus genotypic classes and used the mean phenotypic differences to estimate the effect of allele substitutions.

**Fig. 2.**
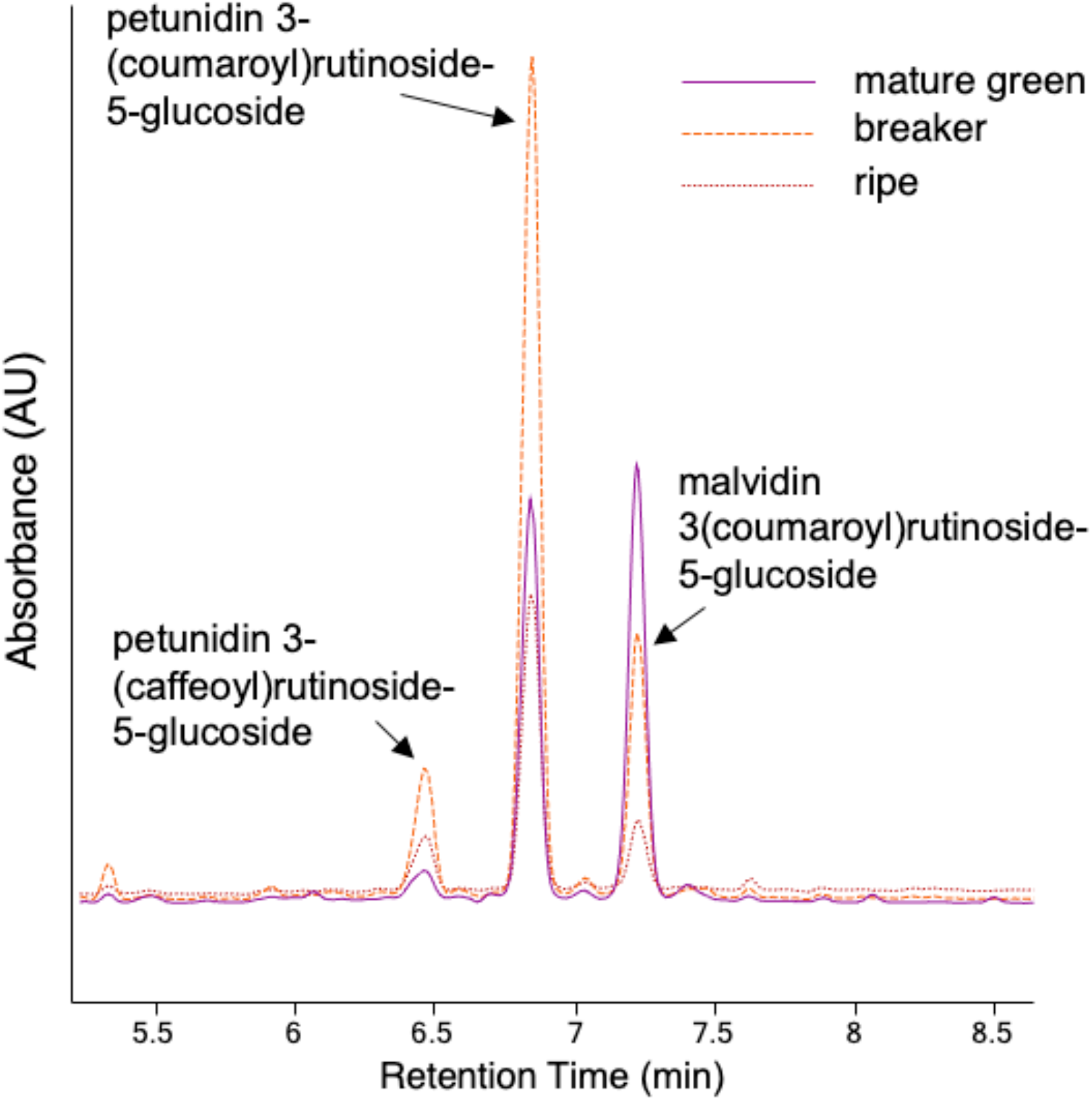
Predominant pigments in the fruit of LA1141 derived lines. The chromatograms show ultra-high performance liquid chromatography separation and photo diode array (UHPLC-PDA) absorbance at 520 nm for fruit from mature green, breaker, and ripe fruit. The predominant peaks were identified as anthocyanins and are labeled above.

Additionally, we tested the pairwise combination of Ant1**_**1 (*Aft*) and atv_ex4 (*atv*), in the IBC and F_2_ populations. We used the linear model *Y* = *μ x* + *M*_*1*_+ *M*_*2*_ + *M*_*1*_:*M*_*2*_ +*E*: where *Y* was the color trait value, *μ x* was equal to the population mean, *M*_*1*_ and M_2_ were effects of individual marker alleles, *M*_*1*_*:M*_*2*_ was the interaction between marker alleles, and *E* was the associated error, equivalent to genotype (marker) to test for significant markers interactions. We conducted a linear model analysis of variance (ANOVA) using the “lm” function in the R core package (R Core Team, 2020) to test the pairwise combination of chromosome 7 (*atv)* and chromosome 10 (*Aft)*. If marker classes were significantly different (*p*<0.05) we used a Tukey’s Honest Significant Difference test, with the “HSD.test” function in the R package Agricolae (De Mendiburu, 2017) to compare means.

### Sequence alignment and phylogeny

A PCR amplification strategy was used for sequencing the *Aft* candidate sequences derived from LA1141 and OH8245. Amplified products were purified by precipitation using a 9:1 ethanol: sodium acetate (3 M), pH 5.2 mixture. Sequencing was performed at the Molecular and Cellular Imaging Center in Wooster, Ohio, using a di-deoxy Sanger procedure on an ABI Prism Sequencer 3100×1 (Grand Island, NY, USA). For each amplicon, the DNA sequence was generated in forward and reverse directions. All sequence data were quality checked and trimmed before alignment. We used the UGENE v. 37 software package (Okonechnikov et al., 2012) to create contigs from the forward and reverse sequence generated sequence from LA1141 and OH8245 corresponding to MYB-encoding genes underling *atv* and *Aft*.

### Bioinformatics pipeline

Genomes from 84 unique cultivated and wild tomato accessions published as part of the 100 Tomato Genome Sequencing Consortium (The 100 Tomato Genome Sequencing Consortium, et al., 2014) and a reference quality whole genome sequence for OH8245 generated as part of a collaboration between the Tomato Pan Genome Consortium and NRGene (Ness-Ziona, Israel; see: https://www.nrgene.com/solutions/consortia/tomato/) were stored on the Ohio Supercomputer Center (OSC) (Ohio Supercomputer center, 1987) computing environment and a nucleotide BLAST database was created using the function “makeblastdb” in the Basic Local Alignment Search Tool (BLAST) version/2020-04 (Altschul et al., 1990) program. Our workflow parsed through sequence matches, identified the highest quality match, and created a FASTA file containing the match as a FASTA output file. Parsing was facilitated by “SearchIO”, “Seq”, and “SeqIO”, functions in BioPerl (Stajich et al., 2002) following implementation of the “blastn” function in the BLAST core package. The steps in the pipeline were automated using the Bash scripting language (Gnu, 2007) in a Unix shell on the OSC.

Passport data for all accessions and a summary of sequences including genomic and coding sequence (CDS) length is available at https://doi.org/10.5281/zenodo.5650141 (Fenstemaker et al., 2021c). The genomic sequences and CDS were retrieved from regions corresponding to the tomato *Aft* locus from LA1141, OH8245, Heinz1706 as described above. CDS corresponding to MYB encoding genes corresponding to LA1141, OH8245 and the 84 tomato accessions published as part of The 100 Tomato Genome Sequencing Consortium (The 100 Tomato Genome Sequencing Consortium, et al., 2014) were determined by comparing genomic sequences to the Heinz reference Tomato Genome CDS (ITAG release 4.0) available from the Solanaceae Genome Network (SGN) (available at: https://solgenomics.net/organism/Solanum_lycopersicum/genome). Additional CDS sequences were retrieved from the National Center for Biotechnology Information (NCBI) from the following Genbank records: Indigo Rose [MN433087 (Yan et al., 2020)], *S. lycopersicum* accession LA1996 [MN242011.1, EF433417.1 (Sapir et al., 2008; Colanero et al., 2020a)], *S. chilense* (Dunal) Reiche (formerly *Lycopersicon chilense* Dunal), and accession LA1930 [MN242012.1 (Colanero et al., 2020a)].

Orthologous sequences corresponding to tomato *Aft* were retrieved from *S. lycopersicoides* Dunal accession LA2951 genome (v0.6) made available by The *Solanum lycopersicoides* Genome Consortium (Powell et al., 2020). *Solanum tuberosum* L. Group Phureja clone DM1-3 genome (PGSC DM v4.03 Pseudomolecules) made available by the Potato Genome Sequence Consortium (PGSC: Potato Genome Sequencing Consortium et al., 2011 and *Capsicum annuum* L., 1753 in [Linnaeus C (1753c)] cv. CM334 genome (Capsicum annuum cv CM334 genome chromosome release 1.55, Hulse-Kemp et al. 2018). These corresponding sequences were retrieved using the BLAST tool at: https://solgenomics.net/tools/blast/. Comparison of syntenic chromosomal regions were made using known marker positions of tomato, potato, and pepper and the comparative map viewer (available at https://solgenomics.net/cview) on chromosome 10. Orthologous sequences corresponding to S*alvia miltiorrhiza* Bunge, 1833 [KF059503.1, (Li and Lu, 2014)], *Arabidopsis thaliana* (L.) Heynh., 1842 (Arabidopsis), [NM_105308.2, NM_105310.4 (Teng et al., 2005, Cominelli et al., 2008; Beradini et al., 2015)] were chosen based on homology and gene annotation that described positive R2R3 MYB regulation of anthocyanins.

The CDS corresponding to the *Aft* genes were retrieved from the CDS reference genomes available from the Sol Genomics Network (SGN): Tomato Genome CDS (ITAG release 4.0), Potato PGSC DM v3.4 CDS sequences, Capsicum annuum cv CM334 Genome CDS (release 1.55) or from NCBI available at: https://www.ncbi.nlm.nih.gov. To retrieve CDS sequence from NCBI, we accessed the “RefSeq” section of the Genbank records mentioned above. The CDS was extracted from the “features” section of the Genbank records and exported as FASTA files. The UGENE v. 37 software package (Okonechnikov et al., 2012) was used for sequence trimming prior to alignment using MUSCLE (version 3.8.31) (Edgar, 2004) in the OSC Unix command line.

Phylogenetic trees were constructed using the phangorn R package (Schliep, 2011) for the R2R3 MYB-encoding genes *Ant1, An2-like*. Genomic sequence files were combined from the MYB encoding genes *An2-like* and *Ant1* to create a single *Aft* locus contig, aligned in MUSCLE and imported into phangorn. We constructed Maximum likelihood trees based on the nucleotide alignment using the general time reversible model with the rate variation among sites described by a gamma distribution and the proportion of invariable sites (a.k.a. GTR+G+I model). The “optim.pml” function was used to optimize model parameters with a stochastic search algorithm to compute the likelihood of the phylogenetic trees (Nguyen et al., 2015). This methodology was used for both genomic and CDS sequences. Clade support was estimated with 1000 bootstrap replicates using the function “bootstrap.pml”. Phylogenetic trees were midpoint rooted for phylogenetic studies that used genomic sequence and rooted using Arabidopsis as an outgroup for phylogenetic studies that used CDS. Trees were drawn and annotated using the Interactive Tree Of Life (ITOL) (Letunic and Bork, 2021; available at https://itol.embl.de/).

## Results

### Accession LA1141 phenotypic description

We observed purple pigmentation in the mature green (MG) fruit of LA1141 (Fig. 1A), and we were able to recover purple fruit in BC_2_S_3_ progenies (Fig. 1B, C). Purple pigmentation occurred in the skin and the pericarp tissues beneath the skin. Pigmentation was visible at all fruit maturity stages, but most apparent at the MG stage. The interior of the fruit did not contain visible purple pigment. Fruit hue values in the inbred backcross (IBC) progenies ranged from 231.27 to 283.35 degrees with a mean of 240.75 degrees for the population. Hue values greater than 250 degrees were designated as “deep purple” (Fig. 1C). Progenies with hue values that ranged between 245 and 250 degrees also had visible spotting or speckling of purple pigment. We designated progenies in this range of hue as “light purple” (Fig. 1B). All hue values measured on fruit below 245 degrees were green (Fig. 1D). Inbred backcross lines (IBLs) with purple pigmentation in the fruit had hue values greater than 245 degrees, L* values ranging from 44.4 to 64.29 units, and chroma values ranging from 7.91 to 35.22 units. We expected the LA1141 × OH8245 BC_2_S_3_ IBC population to be roughly 87.5% recurrent parent (OH8245), with the remaining 12.5% representing random introgressions from the LA1141 donor parent. The observed segregation of individuals with deep purple phenotypes approximated the expected genotypic percentages for two unlinked loci (χ^2^=0.339, *p*=0.843). Plants with deep purple fruit (Fig. 1C) also display darker green leaves with purple veins and purple pigmentation in the stems. In contrast, plants with the light purple phenotypes (Fig.1B) could be explained by a single introgression (χ^2^ =2.053, *p* =0.358).

### Chemical analysis of LA1141 × OH8245 BC_2_S_3_ derived purple tomatoes

We used UHPLC-PDA-QTOF-MS to identify compounds that absorb light at 520 nm, which is characteristic of anthocyanins. The peaks in the chromatogram (Fig. 2) indicate the predominant anthocyanidins were petunidin and malvidin based on accurate masses previously published (Mathews et al., 2003, Ooe et al., 2016). Petunidin-3-(caffeoyl)rutinoside-5-glucoside (C_43_H_49_O_24_ ^+^) was identified at a retention time of 6.46 minutes and had an observed mass [M^+^] of 949.2623 (1 ppm mass error), petunidin-(coumaroyl)rutinoside 5-glucoside (C_43_H_49_O_23_ ^+^) at a retention time of 6.85 minutes with a mass [M^+^] of 933.2686 (2 ppm mass error), and malvidin-3(coumaroyl)rutinoside-5-glucoside (C_44_H_51_O_23_ ^+^) at a retention time of 7.22 minutes with a mass [M^+^] of 947.2834 (1.3 ppm mass error) (Fig. 2). These anthocyanins are present in all fruit maturity stages. We see a change in the predominant anthocyanins from the MG to breaker fruit stage (Fig. 2). The anthocyanins petunidin-(coumaroyl)rutinoside 5-glucoside and anthocyanin malvidin 3(coumaroyl)rutinoside-5-glucoside are of similar intensity at MG (Fig. 2). The anthocyanin Petunidin-(coumaroyl)rutinoside 5-glucoside was the predominant anthocyanin at breaker and ripe stages (Fig. 2). Additionally, we observed changes in individual anthocyanin abundance over ripening (Fig. 2). The anthocyanin Malvidin 3(coumaroyl)rutinoside-5-glucoside was most abundant at the MG stage (Fig. 2). The anthocyanins Petunidin-(coumaroyl)rutinoside 5-glucoside and petunidin-3-(caffeoyl)rutinoside-5-glucoside are most abundant at the breaker stage (Fig. 2).

### LA1141 × OH8245 BC_2_S_3_ linkage map quality

Linkage maps were constructed based on marker data from the BC_2_S_3_ IBC population and defined 13 linkage groups corresponding to each tomato chromosome. We split chromosome 1 into two linkage groups (1a and 1b) because of a recombination fraction greater than 0.45 between adjacent markers. The total number of markers in each linkage group ranged between 2 and 27, and linkage group 4 had the most markers at 27 (Table 1). The centimorgan (cM) length per linkage group ranged between 25.8 and 121.6 cM (Table 1). The average cM distance between markers was 8.1, and the largest distance in cM between markers was 41.8 (Table 1). Single nucleotide polymorphism (SNP) marker physical position using the tomato Sl4.0 physical map (Hosmani et al., 2019) agreed with the estimated genetic position using both linear correlation and rank correlation (Table 1). As previously demonstrated, correlations are not perfectly linear due to reduced recombination in the centromere (Sim et al., 2012). Linear correlations ranged from 0.28-0.99, while rank correlations ranged from 0.96 to 1 (Table 1).

**Table 1.**
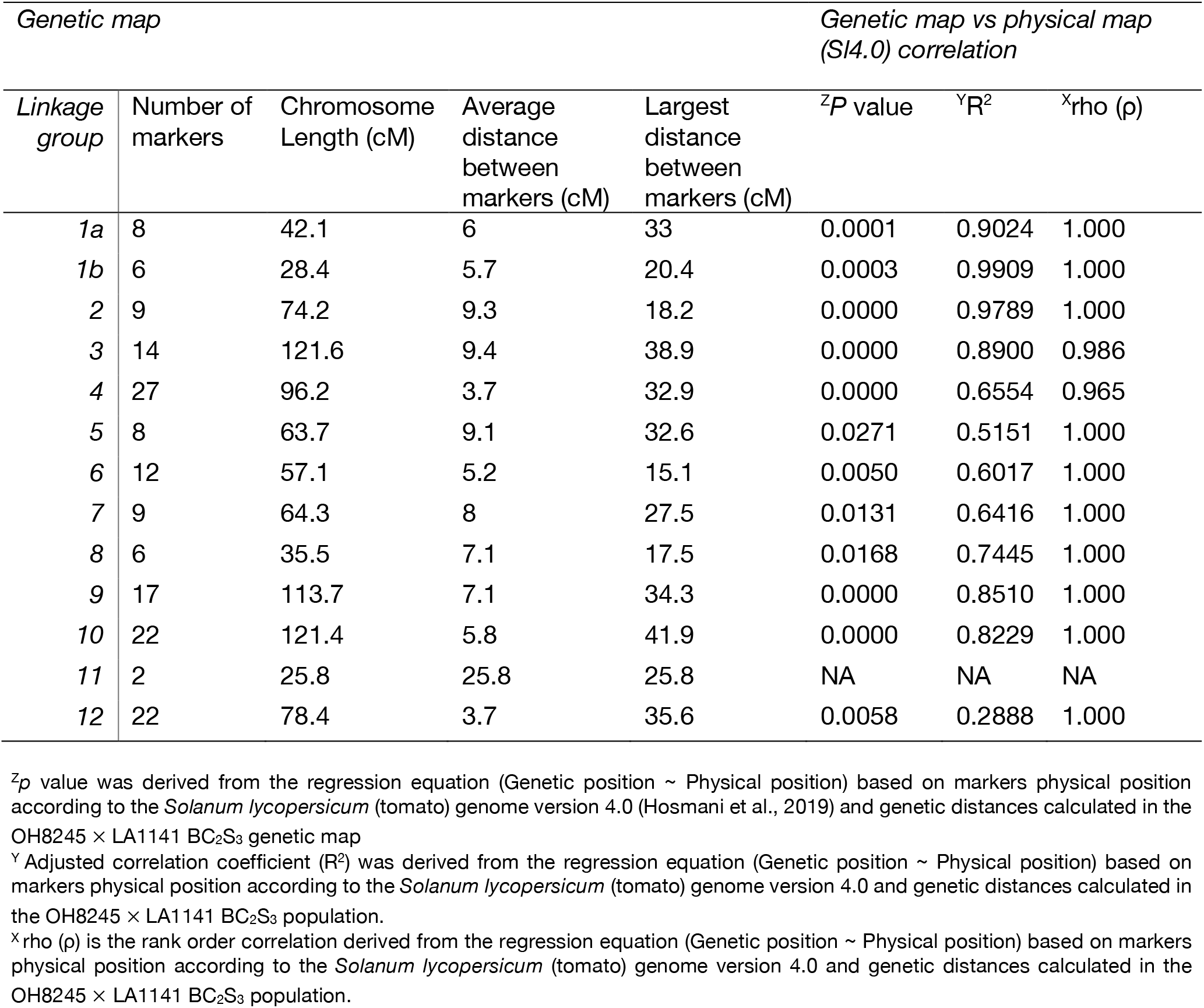
Genetic map quality for the inbred backcross population (OH8245 × LA1141 BC_2_S_3_).

| *Genetic map* |  |  |  |  | *Genetic map vs physical map (SI4.0) correlation* |  |  |
| --- | --- | --- | --- | --- | --- | --- | --- |
| *Linkage group* | Number of markers | Chromosome Length (cM) | Average distance between markers (cM) | Largest distance between markers (cM) | <sup>z</sup>P value | <sup>y</sup>R<sup>2</sup> | <sup>x</sup>rho (ρ) |
| 1a | 8 | 42.1 | 6 | 33 | 0.0001 | 0.9024 | 1.000 |
| 1b | 6 | 28.4 | 5.7 | 20.4 | 0.0003 | 0.9909 | 1.000 |
| 2 | 9 | 74.2 | 9.3 | 18.2 | 0.0000 | 0.9789 | 1.000 |
| 3 | 14 | 121.6 | 9.4 | 38.9 | 0.0000 | 0.8900 | 0.986 |
| 4 | 27 | 96.2 | 3.7 | 32.9 | 0.0000 | 0.6554 | 0.965 |
| 5 | 8 | 63.7 | 9.1 | 32.6 | 0.0271 | 0.5151 | 1.000 |
| 6 | 12 | 57.1 | 5.2 | 15.1 | 0.0050 | 0.6017 | 1.000 |
| 7 | 9 | 64.3 | 8 | 27.5 | 0.0131 | 0.6416 | 1.000 |
| 8 | 6 | 35.5 | 7.1 | 17.5 | 0.0168 | 0.7445 | 1.000 |
| 9 | 17 | 113.7 | 7.1 | 34.3 | 0.0000 | 0.8510 | 1.000 |
| 10 | 22 | 121.4 | 5.8 | 41.9 | 0.0000 | 0.8229 | 1.000 |
| 11 | 2 | 25.8 | 25.8 | 25.8 | NA | NA | NA |
| 12 | 22 | 78.4 | 3.7 | 35.6 | 0.0058 | 0.2888 | 1.000 |

<sup>z</sup>p value was derived from the regression equation (Genetic position ~ Physical position) based on markers physical position according to the *Solanum lycopersicum* (tomato) genome version 4.0 (Hosmani et al., 2019) and genetic distances calculated in the OH8245 × LA1141 BC<sub>2</sub>S<sub>3</sub> genetic map
<sup>y</sup> Adjusted correlation coefficient (R<sup>2</sup>) was derived from the regression equation (Genetic position ~ Physical position) based on markers physical position according to the *Solanum lycopersicum* (tomato) genome version 4.0 and genetic distances calculated in the OH8245 × LA1141 BC<sub>2</sub>S<sub>3</sub> population.
<sup>x</sup> rho (ρ) is the rank order correlation derived from the regression equation (Genetic position ~ Physical position) based on markers physical position according to the *Solanum lycopersicum* (tomato) genome version 4.0 and genetic distances calculated in the OH8245 × LA1141 BC<sub>2</sub>S<sub>3</sub> population.

### Quantitative trait loci analysis of tomato color in the LA1141× OH8245 BC_2_S_3_ population

We identified three putative QTLs that explained between 8.24 and 35.53% of the total phenotypic variation for hue, chroma and L* (Fig. 3; Table 2). All QTLs that contribute to purple color are derived from the LA1141 donor parent with purple pigmentation defined by an increase in hue and a decrease in both chroma and L* (Table 2). A region on the distal arm of chromosome 10 explained between 22.63 and 24.04% of the total phenotypic variation, and increased hue between 6.74 and 7.5 degrees (Fig. 3; Table 2). Two QTLs, one on the proximal arm and one on the distal arm of chromosome 10, were associated with chroma and explained between 18.02 and 28.53% (proximal arm) and between 15.95 and 23.08% (distal arm) of the total phenotypic variance (Fig. 3; Table 2). These QTLs decreased chroma between 3.96 and 17.53 units (proximal arm) and 6.26 and 8.42 units (distal arm) (Fig. 3; Table 2). Two QTLs were associated with L*, one on chromosome 6 and one on chromosome 10 (proximal) (Fig. 3; Table 2). These QTLs explained between 8.24 and 35.53% of the total phenotypic variation (Table 2). The QTL on chromosome 10 explained between 22.03 and 35.53% of the phenotypic variation and reduced L* by 9.23 units (Table 2). The QTL on chromosome 6 explained between 8.24 and 10.13% of the phenotypic variation and reduced L* between 4.53 and 5.05 units (Table 2).

**Table 2.**
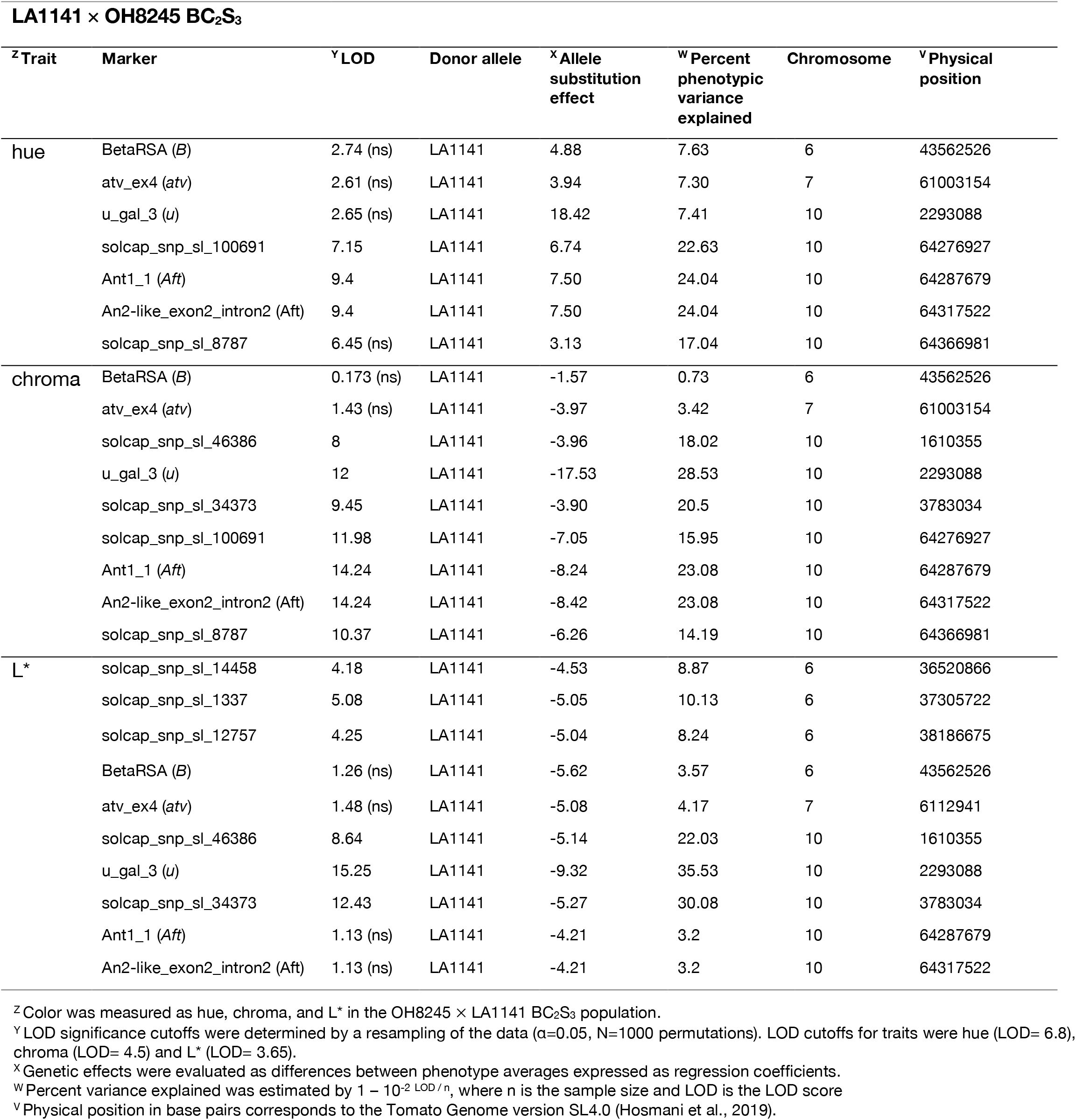
Markers associated with tomato fruit color.

| **LA1141 × OH8245 BC<sub>2</sub>S<sub>3</sub>** |  |  |  |  |  |  |  |
| --- | --- | --- | --- | --- | --- | --- | --- |
| <sup>z</sup>Trait | Marker | <sup>y</sup>LOD | Donor allele | <sup>x</sup>Allele substitution effect | <sup>w</sup>Percent phenotypic variance explained | Chromosome | <sup>v</sup>Physical position |
| hue | BetaRSA (*B*) | 2.74 (ns) | LA1141 | 4.88 | 7.63 | 6 | 43562526 |
|  | atv_ex4 (*atv*) | 2.61 (ns) | LA1141 | 3.94 | 7.30 | 7 | 61003154 |
|  | u_gal_3 (*u*) | 2.65 (ns) | LA1141 | 18.42 | 7.41 | 10 | 2293088 |
|  | solcap_snp_sl_100691 | 7.15 | LA1141 | 6.74 | 22.63 | 10 | 64276927 |
|  | Ant1_1 (*Aft*) | 9.4 | LA1141 | 7.50 | 24.04 | 10 | 64287679 |
|  | An2-like_exon2_intron2 (*Aft*) | 9.4 | LA1141 | 7.50 | 24.04 | 10 | 64317522 |
|  | solcap_snp_sl_8787 | 6.45 (ns) | LA1141 | 3.13 | 17.04 | 10 | 64366981 |
| chroma | BetaRSA (*B*) | 0.173 (ns) | LA1141 | -1.57 | 0.73 | 6 | 43562526 |
|  | atv_ex4 (*atv*) | 1.43 (ns) | LA1141 | -3.97 | 3.42 | 7 | 61003154 |
|  | solcap_snp_sl_46386 | 8 | LA1141 | -3.96 | 18.02 | 10 | 1610355 |
|  | u_gal_3 (*u*) | 12 | LA1141 | -17.53 | 28.53 | 10 | 2293088 |
|  | solcap_snp_sl_34373 | 9.45 | LA1141 | -3.90 | 20.5 | 10 | 3783034 |
|  | solcap_snp_sl_100691 | 11.98 | LA1141 | -7.05 | 15.95 | 10 | 64276927 |
|  | Ant1_1 (*Aft*) | 14.24 | LA1141 | -8.24 | 23.08 | 10 | 64287679 |
|  | An2-like_exon2_intron2 (*Aft*) | 14.24 | LA1141 | -8.42 | 23.08 | 10 | 64317522 |
|  | solcap_snp_sl_8787 | 10.37 | LA1141 | -6.26 | 14.19 | 10 | 64366981 |
| L* | solcap_snp_sl_14458 | 4.18 | LA1141 | -4.53 | 8.87 | 6 | 36520866 |
|  | solcap_snp_sl_1337 | 5.08 | LA1141 | -5.05 | 10.13 | 6 | 37305722 |
|  | solcap_snp_sl_12757 | 4.25 | LA1141 | -5.04 | 8.24 | 6 | 38186675 |
|  | BetaRSA (*B*) | 1.26 (ns) | LA1141 | -5.62 | 3.57 | 6 | 43562526 |
|  | atv_ex4 (*atv*) | 1.48 (ns) | LA1141 | -5.08 | 4.17 | 7 | 6112941 |
|  | solcap_snp_sl_46386 | 8.64 | LA1141 | -5.14 | 22.03 | 10 | 1610355 |
|  | u_gal_3 (*u*) | 15.25 | LA1141 | -9.32 | 35.53 | 10 | 2293088 |
|  | solcap_snp_sl_34373 | 12.43 | LA1141 | -5.27 | 30.08 | 10 | 3783034 |
|  | Ant1_1 (*Aft*) | 1.13 (ns) | LA1141 | -4.21 | 3.2 | 10 | 64287679 |
|  | An2-like_exon2_intron2 (*Aft*) | 1.13 (ns) | LA1141 | -4.21 | 3.2 | 10 | 64317522 |

<sup>z</sup> Color was measured as hue, chroma, and L\* in the OH8245 × LA1141 BC<sub>2</sub>S<sub>3</sub> population.
<sup>y</sup> LOD significance cutoffs were determined by a resampling of the data ( $\alpha=0.05$ , $N=1000$ permutations). LOD cutoffs for traits were hue (LOD= 6.8), chroma (LOD= 4.5) and L\* (LOD= 3.65).
<sup>x</sup> Genetic effects were evaluated as differences between phenotype averages expressed as regression coefficients.
<sup>w</sup> Percent variance explained was estimated by $1 - 10^{-2 \text{ LOD} / n}$ , where $n$ is the sample size and LOD is the LOD score
<sup>v</sup> Physical position in base pairs corresponds to the Tomato Genome version SL4.0 (Hosmani et al., 2019).

**Fig. 3.**
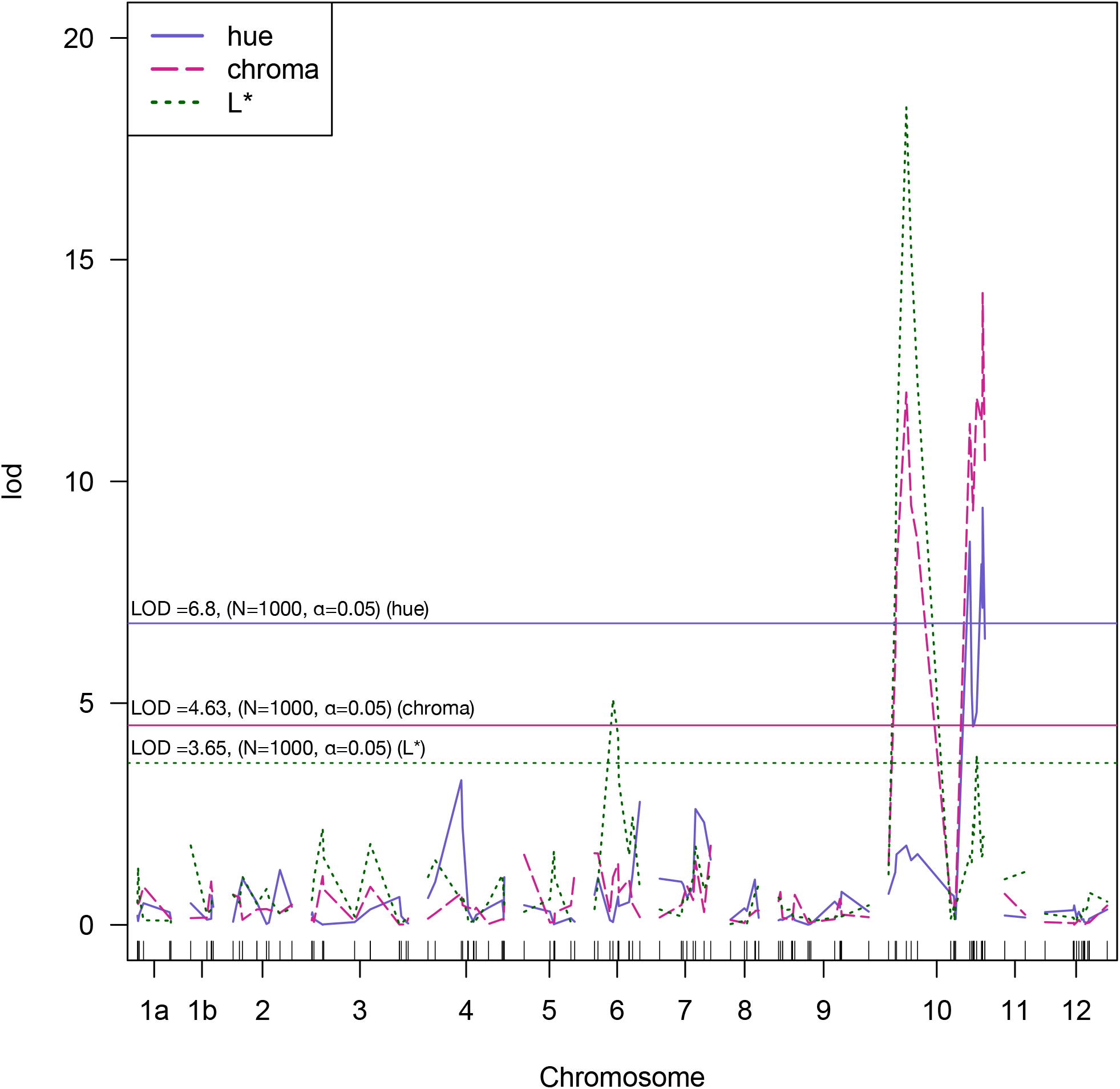
Composite interval mapping (CIM) of fruit color measured as hue (violet), chroma (pink), and L* (green, dotted) in the LA1141 x OH8245 BC_2_S_3_ inbred backcross population. The y-axis is the logarithm of the odds (LOD). The horizontal lines are the resampled LOD significance cutoff (α=0.05, N=1000 permutations) for hue (violet), chroma (pink) and L* (green, dotted). The x-axis represents the 12 chromosomes in tomato and chromosome distance in cM was calculated using the Kosambi function to correct for multiple crossovers.

### Candidate genes

Candidate genes were selected because of their previously characterized role in regulating tomato fruit pigmentation and because of their locations within the physical interval of our QTL (Table 2). The R2R3 MYB-encoding candidate genes *Ant1 (Aft)* (Sapir et al., 2008) and *An2-like* (*Aft*) (Qiu et al., 2019; Yan et al., 2020) are located within the QTL interval on the distal arm of chromosome 10 (Table 2). The MYB encoding genes *Ant1* and *An2-like* are members of the multi-gene MYB family associated with the *Aft* locus (Yan et al., 2020). The transcription factor *Golden2-like 2* (*u*) (Powell et al., 2012) maps to the proximal arm of chromosome 10 within the QTL regions identified for L* and chroma (Table 2). Additionally, we chose the fruit-specific *Cyc-B* gene *(B)* to investigate the QTL on chromosome 6 because accession LA1141 has the characteristic ripe orange fruit associated with the *Beta* locus (Orchard et al., 2021). We chose The R3 MYB repressor *atv* (*atroviolacea*) on chromosome 7 (Cao et al., 2017; Colanero et al., 2018) because of its previously described synergistic interaction with *Aft* which results in a purple phenotype similar to what we observe in our deep purple accession (Fig. 1C). We added these markers to the linkage maps described above and used them in our QTL analysis.

### QTL mapping using candidate genes in the IBC population

Genetic evidence supports a role for *Aft, atv* and *u* conferring purple pigmentation in the fruit of LA1141. The markers corresponding to the MYB-encoding genes *Ant1* (Ant1_1 *(Aft*)) and *An2-like* (An2-like_exon2_intron2 (*Aft*)) are physically near one another (Hosmani et al., 2019) (Table 2) and genetically linked (χ2 =3.36, *p*=0.186). For measurements of hue, the markers Ant1_1 (*Aft*) and An2-like_exon2_intron2 (*Aft*) (LOD=9.4) fell above our resampled logarithm of the odds (LOD) cutoff (LOD=6.8), explained 24.04 % of the phenotypic variation, and increased hue by 7.05 degrees (Table 2). The markers BetaRSA (*B*) (LOD=2.74), atv_ex4 (*atv*) (LOD=2.6), and u_gal_3 (*u*) (LOD=2.65) did not fall above our resampled LOD cutoffs for hue (Table 2).

The markers Ant1_1 (*Aft*) and An2-like_exon2_intron2 (*Aft*) (LOD=14.24) fell above our resampled LOD cutoffs for chroma (LOD=4.5). The markers Ant1_1 (*Aft*) and An2-like_exon2_intron2 (*Aft*) explained 23.08% of the total phenotypic variance and reduced chroma by 8.24 units (Table 2). The marker u_gal_3 (*u*) (LOD=12) also fell above our resampled LOD cutoffs, explained 28.53 % of the total phenotypic variation, and reduced chroma by 17.53 units (Table 2). The markers BetaRSA (*B*) (LOD=2.74) and atv_ex4 (*atv*) (LOD=2.61) did not fall above our resampled LOD cutoff for chroma (Table 2).

Regions on chromosome 6 and the proximal arm of chromosome 10 were targeted for measurements of L*. The marker u_gal_3 (*u*) (LOD=15.25) fell above our resampled LOD cutoff (LOD=3.65). The marker u_gal_3 (*u*) explained 35.53 % of the total phenotypic variance and reduced L* by 9.32 units (Table 2). The marker BetaRSA (*B*) (LOD=1.26) did not fall above our resampled LOD cutoff for L* and our QTL analysis failed to support a role for *B* as a candidate gene on chromosome 6. Additionally, the markers Ant1_1 (*Aft*) and An2-like_exon2_intron2 (*Aft*) (LOD=1.13), and atv_ex4 (*atv*) (LOD=1.48) did not appear to be associated with L* (Table 2).

Although the marker atv_ex4 (*atv*) did not fall above our LOD significance thresholds for hue, chroma, or L* (Table 2), segregation rates of the deep purple phenotype in the BC_2_S_3_ progenies suggested two unlinked loci were responsible. The known regulatory mechanism involving MYB encoding genes underlying *atv* and *Aft* led us to pursue the interaction effects of the combined loci on chromosome 7 and on chromosome 10. The interaction between homozygous LA1141 *Aft* (*Aft*/*Aft)* and the homozygous LA1141 *atv* (*atv/atv*) was significant (*p*=< 2.2e-16) (Fig. 4). We compared the hue values of BC_2_S_3_ IBL progenies that were *Aft/Aft atv/atv* to homozygous OH8245 *Aft* (*AFT*/*AFT*) and homozygous OH8245 *atv* (*ATV/ATV*) (Fig. 4A). The BC_2_S_3_ IBLs with both the *Aft* and *atv* locus, which is notated as the *Aft/Aft atv/atv* genotype had higher hue values than the *AFT/AFT ATV/ATV* genotypes (Fig. 4A). Additionally, we compared all possible marker-locus class combinations, including the genotypes *Aft/Aft ATV/ATV*, and *AFT/AFT atv/atv*. The *Aft/Aft atv/atv* genotype had higher hue values than all other genotypes (Fig. 4A). However, the *Aft/Aft ATV/ATV* genotype had higher hue values than the *AFT/AFT atv/atv* and *AFT/AFT ATV/ATV* genotypes (Fig. 4A).

**Fig. 4.**
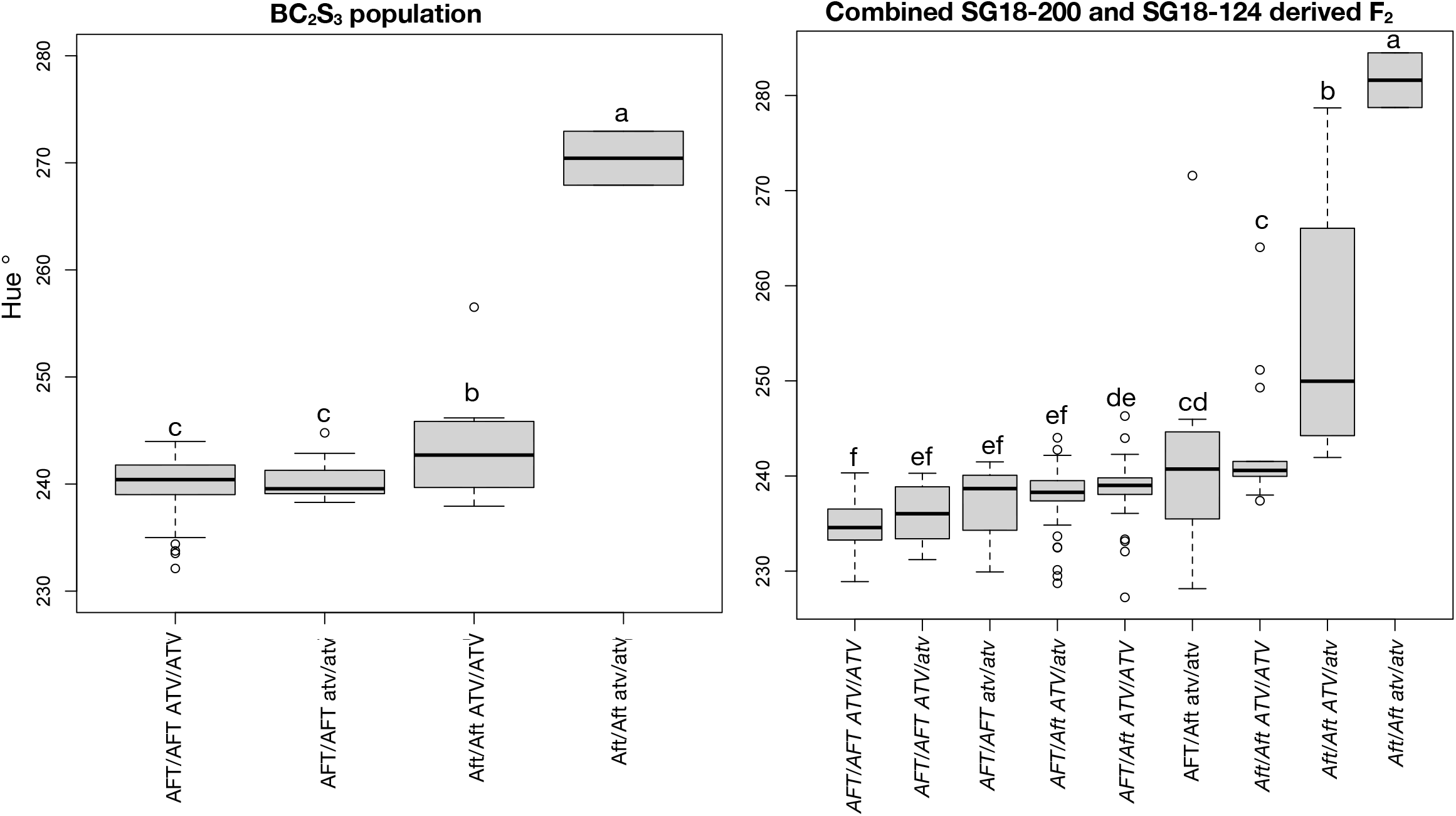
Box plots represent interactions between the *Anthocyanin fruit* and *atroviolacium* loci. The x-axis is marker-locus genotypic class, and the y-axis is degrees of hue. **(A)** The interaction is shown in the BC_2_S_3_ population and (**B)** the combined F_2_ validation populations. For the *Anthocyanin fruit* locus: homozygous LA1141 alleles are abbreviated as *Aft/Aft*, heterozygous alleles as *AFT/aft*, and homozygous OH8245 *AFT*/*AFT*. For the *atroviolacium* locus: homozygous LA1141 alleles are abbreviated as *atv/atv*, heterozygous alleles as *ATV/atv*, and homozygous OH8245 *ATV/ATV*. Different letters indicate statistically significant differences among groups (Tukey’s Honest Significant Difference (HSD), P<0.05). Marker-locus genotypic class notation follows previous pblications (Cao at al. 2017).

### Confirmation of QTLs in the F_2_ validation populations

We evaluated F_2_ populations originating from the selected IBL progenies SG18-124 (Fig. 1C) and SG18-200 (Fig. 1B) for measurements of hue, chroma, and L* to validate the QTLs identified in the BC_2_S_3_ generation. The IBL SG18-124 had deep purple fruit (Fig. 1C). The mean hue value of the SG18-124 derived F_2_ population was 238.5 degrees and ranged from 227.24 to 284.4 degrees. The mean chroma value was 24.8 units and ranged from 5.7 to 39 units. The mean L* value was 46.3 units and ranged from 30.3 to 67.1 units. The IBL SG18-200 had light purple fruit (Fig. 1B). The mean hue value in the SG18-200 derived F_2_ population was 239.7 degrees and ranged from 234.8 to 264 degrees. The mean chroma value was 29.1 and ranged from 13.7 to 33.79 units. The mean L* value was 52.2 and ranged from 42.3 to 60.2 units.

In the SG18-124 derived F_2_ population the markers Ant1_1 (*Aft*) (P=1.513e-09), An2-like_exon2_intron2 (*Aft*) (*p*=2.118e-09) were significantly associated with hue. The markers Ant1_1 (*Aft*) and An2-like_exon2_intron2 (*Aft*) both explained 37% of the phenotypic variation, and increased hue by 19.45 and 22.05 degrees respectively (Table 3). The marker atv_ex4 (*atv*) (*p*=0.022) was also significantly associated with hue, explained 9% of the phenotypic variation, and increased hue by 11.99 degrees (Table 3). The marker u_gal_3 (*u*) (*p*=0.901) was not significant for hue (Table 3). However, the marker u_gal_3 (*u*) (*p*=2.071e-05) was significantly associated with chroma, explained 23% of the phenotypic variation, and decreased chroma by 10.67 units (Table 3). The markers An2-like_exon2_intron2 (*Aft*) (*p*=4.051e-04 and Ant1_1 (*Aft*) (*p*=9.009e-07) were significantly associated with chroma, explained 14% and 27% of the total phenotypic variation, and decreased chroma by 10.80 and 12.23 units (Table 3). The marker u_gal_3 (*u*) (*p* = 3.181e-04) was significantly associated with L*, explained 17 % of the phenotypic variation, and decreased L* by 10.38 units (Table 3). The marker BetaRSA (*B*) was not significantly associated with hue (*p*=0.103), chroma (*p*=0.842), or L* (*p*=0.715) in the SG18-124 derived F_2_ population (Table 3).

**Table 3.** Candidate gene associations validated in subsequent F_2_ populations.

| **SG18-124 IBL derived F<sub>2</sub> validation population** |  |  |  |  |  |  |  |
| --- | --- | --- | --- | --- | --- | --- | --- |
| <sup>z</sup>Trait | Marker | <sup>y</sup>P value | Parent allele | <sup>x</sup>Allele substitution effect | <sup>w</sup>R<sup>2</sup> | Chromosome | <sup>v</sup>Physical position |
| hue | BetaRSA (*B*) | 0.103 (ns) | LA1141 | -0.63 | 0.04 | 6 | 43562526 |
|  | atv_ex4 (*atv*) | 0.022 | LA1141 | 11.99 | 0.09 | 7 | 61003154 |
|  | u_gal_3 (*u*) | .901 (ns) | LA1141 | 0.4026 | -0.02 | 10 | 2293088 |
|  | Ant1_1 (*Aft*) | <0.000 | LA1141 | 19.45 | 0.37 | 10 | 64287679 |
|  | An2-like_exon2_intron2 (*Aft*) | <0.000 | LA1141 | 22.05 | 0.37 | 10 | 64366981 |
| Chroma | BetaRSA (*B*) | 0.842 (ns) | LA1141 | -1.47 | -0.02 | 6 | 43562526 |
|  | atv_ex4 (*atv*) | 0.06 (ns) | LA1141 | -6.62 | 0.06 | 7 | 61003154 |
|  | u_gal_3 (*u*) | <0.000 | LA1141 | -10.67 | 0.23 | 10 | 2293088 |
|  | Ant1_1 (*Aft*) | <0.000 | LA1141 | -12.23 | 0.27 | 10 | 64287679 |
|  | An2-like_exon2_intron2 (*Aft*) | <0.000 | LA1141 | -10.80 | 0.14 | 10 | 64366981 |
| L* | BetaRSA (*B*) | 0.715 (ns) | LA1141 | -2.41 | -0.02 | 6 | 43562526 |
|  | atv_ex4 (*atv*) | 0.452 (ns) | LA1141 | -3.65 | -0.01 | 7 | 61003154 |
|  | u_gal_3 (*u*) | <0.000 | LA1141 | -10.38 | 0.17 | 10 | 2293088 |
|  | Ant1_1 (*Aft*) | 0.136 (ns) | LA1141 | -4.24 | 0.02 | 10 | 64287679 |
|  | An2-like_exon2_intron2 (*Aft*) | 0.1876 (ns) | LA1141 | -4.55 | 0.01 | 10 | 64366981 |
| **SG18-200 IBL derived F<sub>2</sub> validation population** |  |  |  |  |  |  |  |
| hue | BetaRSA (*B*) | 0.001 | LA1141 | 3.23 | 0.14 | 6 | 43562526 |
|  | atv_ex4 (*atv*) (monomorphic) | NA | OH8245 | NA | NA | 7 | 61003154 |
|  | u_gal_3 (*u*) (monomorphic) | NA | OH8245 | NA | NA | 10 | 2293088 |
|  | Ant1_1 (*Aft*) | <0.000 | LA1141 | 4.36 | 0.17 | 10 | 64287679 |
|  | An2-like_exon2_intron2 (*Aft*) | <0.000 | LA1141 | 5.03 | 0.23 | 10 | 64366981 |
| chroma | BetaRSA (*B*) | 0.06 (ns) | LA1141 | -2.53 | 0.04 | 6 | 43562526 |
|  | atv_ex4 (*atv*) (monomorphic) | NA | OH8245 | NA | NA | 7 | 61003154 |
|  | u_gal_3 (*u*) (monomorphic) | NA | OH8245 | NA | NA | 10 | 2293088 |
|  | Ant1_1 (*Aft*) | <0.000 | LA1141 | -9.04 | 0.48 | 10 | 64287679 |
|  | An2-like_exon2_intron2 (*Aft*) | <0.000 | LA1141 | -9.15 | 0.52 | 10 | 64366981 |
| L* | BetaRSA (*B*) | 0.186 (ns) | LA1141 | -1.97 | 0.01 | 6 | 43562526 |
|  | atv_ex4 (*atv*) (monomorphic) | NA | OH8245 | NA | NA | 7 | 61003154 |
|  | u_gal_3 (*u*) (monomorphic) | NA | OH8245 | NA | NA | 10 | 2293088 |
|  | Ant1_1 (*Aft*) | <0.000 | LA1141 | -6.15 | 0.24 | 10 | 64287679 |
|  | An2-like_exon2_intron2 (*Aft*) | <0.000 | LA1141 | -5.75 | 0.25 | 10 | 64366981 |

<sup>z</sup> Color was measured as hue, chroma, and L\* in the BC<sub>2</sub>S<sub>3</sub> IBL derived F<sub>2</sub> populations.
<sup>y</sup> ANOVAs were conducted, and F-tests were used to determine if significant variation in hue, chroma, and L\* was associated with differences in marker-locus genotypic classes. If NA, the marker was not segregating in the population and therefore could not be tested for differences in marker-locus genotypic classes.
<sup>x</sup> F-tests to determine if hue, chroma, and L\* were associated with significant differences in marker-locus genotypic classes and used the line mean differences to estimate the effect of allele substitutions.
<sup>w</sup> Adjusted correlation coefficient (R<sup>2</sup>) calculated from linear model analysis of variance (ANOVA) is the percent of total phenotypic variance explained.
<sup>v</sup> Physical position in base pairs corresponds to the Tomato Genome version SL4.0 (Hosmani et al., 2019).

In the SG18-200 derived F_2_ population, the markers atv_ex4 *(atv*) and u_gal_3 (*u*) were monomorphic (Table 3). Therefore, we did not test the estimated effects of allele substitutions and associations in this population. The markers Ant1_1 (*Aft*) (*p*=5.702e-04) and An2-like_exon2_intron2 (*Aft*) (*p*=3.691e-05) were significantly associated with hue (Table 3). The markers Ant1_1 (*Aft*) and An2-like_exon2_intron2 (*Aft*) explained 17 and 23% of the phenotypic variation, and increased hue by 4.36 and 5.03 degrees (Table 3). Although the marker BetaRSA (*B*) was not significantly associated with hue in the SG18-124 derived F_2_ population described above, it was significantly associated with the SG18-200 population (*p*=0.001) (Table 3). The marker BetaRSA (*B*) explained 14% of the total phenotypic variation and increased hue by 3.23 degrees (Table 3). The markers Ant1_1 (*Aft*) (*p*=1.475e-09) and An2-like_exon2_intron2 (*Aft*) (*p*= 7.13e-11) were significantly associated with chroma, explained 48% and 52% of the phenotypic variation, and decreased chroma by 9.04 and 9.15 units (Table 3). The marker BetaRSA (*B*) (*p*=0.06) was marginally non-significant for chroma (Table 3). The markers Ant1_1 (*Aft*) (*p*= 7.042e-05) and An2-like_exon2_intron2 (*Aft*) (*p*= 2.296e-05) were significantly associated with L*, explained 24% and 25% of the total phenotypic variation, and decreased L* by 6.15 and 5.75 units (Table 3).

We validated the interaction between homozygous *Aft* (*Aft/Aft*) and homozygous *atv* (*atv/atv*) in the F_2_ progeny (Fig. 4B). Our results confirm an interaction between *Aft* and *atv* is needed for the deep purple fruit phenotype (Fig. 1C) and a single introgression of *Aft* confers purple pigmentation, designated as a light purple phenotype (Fig. 1B). Progeny homozygous for *Aft/Aft atv/atv* genotypes had higher hue values compared to all other marker-locus classes (Fig. 4B). Homozygous *Aft* (*Aft/Aft*) and heterozygous *atv* (*ATV/atv*) also had higher hue values than other marker locus classes, except for the *Aft/Aft atv/atv* genotype (Fig. 4B). These results suggested that the heterozygous *atv* genotype can accumulate enough anthocyanins to measure differences in hue. The *Aft/Aft ATV/ATV* and *AFT/Aft atv/atv* genotypes had higher degrees of hue than the *AFT/AFT ATV/ATV, AFT/AFT atv/atv, AFT/AFT, ATV/atv*, and *AFT/Aft ATV/atv* genotypes (Fig. 4B). Still, they had significantly lower hue values than the *Aft/Aft atv/atv* and *Aft/Aft ATV/atv* genotypes (Fig. 4B).

### Sequence analysis of candidate genes

Sequence reads for *atv* covered 1353 bps (100%) from the first putative start codon. Sequence analysis suggested that the LA1141 *atv* may be nonfunctional compared to the cultivated accessions OH8245 and Heinz 1706. There is an 18 bp INDEL in the first intron of the LA1141 *atv* sequence and two G to A SNPs in the coding region of the second exon (Fig. 5). These G to A SNPs in the coding region may result in the loss of a functional R3/bHLH binding domain (Fig. 5). The LA1141 *atv* sequence is distinct from the allele previously described in Indigo Rose derived from *S. cheesmaniae* accession LA0434 and does not have the previously chracterized 4 bp TAGA insertion (Fig. 5).

**Fig. 5.**
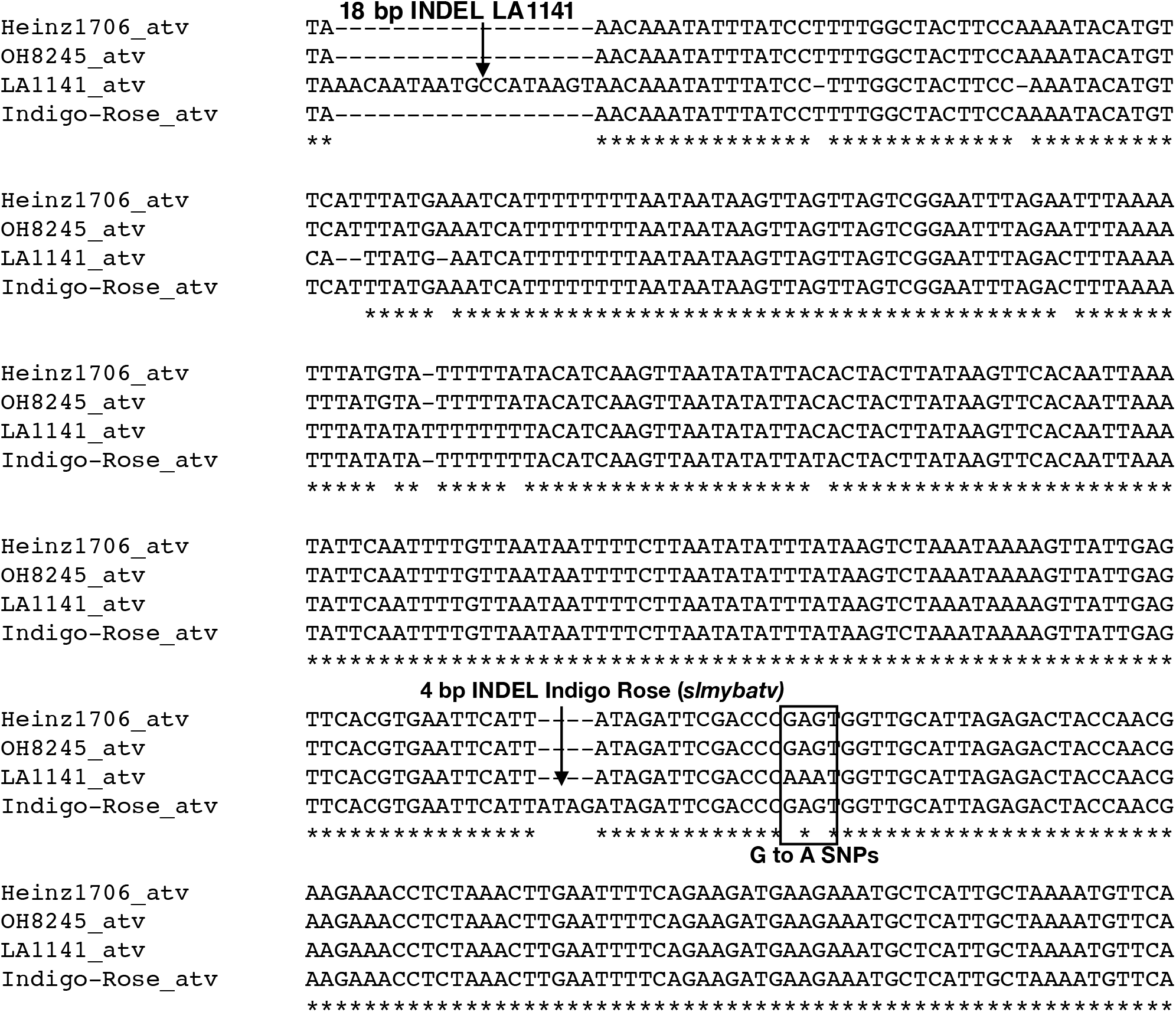
Sequence polymorphism of selected genomic regions of the *atv* locus. A novel 18 bp insertion/deletion (INDEL) found in the first intron in LA1141 is highlighted (arrow). The causal 4 bp INDEL (*slmybatv*) previously characterized in the tomato cultivar Indigo Rose (Cao et al., 2017) is also highlighted (arrow). Two G to A SNPs in the coding region of the 2^nd^ exon (**boxed**) are labeled (**bold**) in a region identified by CRISPR/CAS9 as important for the function of the conserved R3 domain (Yan et al., 2020). Sequences were aligned using MUSCLE (Edgar, 2004) using default settings. Conserved nucleotides are starred. The Heinz reference sequence (Heinz1706), OH8245, LA1141 and Indigo Rose genomic *atv* sequences are represented.

Contigs assembled from sequencing reads of the LA1141 and OH8245 of R2R3 MYB-encoding gene *An2-like* covered approximately 1,363 out of 1,356 base pairs (bps) from the putative start codon. FASTA files corresponding to sequences for tomato accessions used in this study are available at: https://doi.org/10.5281/zenodo.5649546 for *An2-like* and https://doi.org/10.5281/zenodo.5649996 for *Ant1* (Fenstemaker et al., 2021d, e). There were several unique SNPs and INDELs in the LA1141 *An2-like* sequence but none of them were in the conserved R2R3 domains (Fenstemaker et al., 2021d) However, LA1141 possess the previously characterized G to A SNP in the 5’ splice site of the 2nd intron (Sun et al., 2020; Yan et al., 2020; Fenstemaker et al., 2021d). Sequencing reads covered 1182 out of 1012 of LA1141 and 1012 out of 1012 bps of OH8245 from the first putative start codon in the R2R3 MYB-encoding gene *Ant1*. In the 3rd exon of the LA1141 *Ant1* sequence, there is 170 bp insertion/deletion (INDEL) which contained MYB core type 1 and type 2 cis-regulatory elements, an AC rich sequence type 2 cis-regulatory element (Fenstemaker et al., 2021e). Sequence analysis suggests that LA1141 may have a functional R2R3 MYB activator at *Aft* and the R3 MYB repressor corresponding to *atv* is likely nonfunctional. Additional characterization of transcripts, proteins, and protein interactions are needed for *An2-like, Ant1* and *atv* for the confirmation of functional changes.

### Phylogenetic analysis of Aft

We combined the genomic sequences from LA1141, OH8245, and 84 re-sequenced accessions representative of the *Lycopersicon, Arcanum, Eriopersicon*, and *Neolycopersicon* groups. The red-fruited clade is represented by commercial, landrace, and heirloom tomato varieties, and *S. lycopersicum* cerasiforme. This clade also includes *S. pimpinellifolium* and the orange-fruited Galápagos species *S. cheesmaniae* and *S. galapagense*. The green-fruited clade is represented by *Solanum arcanum, S. chilense, S. chmielewskii, S. habrochaites, S. huaylasense, S. neorickii, S. pennellii*, and *S. peruvianum*. Genomic sequence corresponding to *Ant1* ranged from 1023 to 1993 bps and genomic sequences corresponding to *An2-like* ranged from 2292 to 2547 bps (Fenstemaker et., 2021c). The differences in contig length correspond to insertions and deletions within the sequences as contigs matched at the 5’ and 3’ ends.

The maximum likelihood (ML) model phylogeny of the R2R3 MYBs representing *Aft* (Fenstemaker et al., 2021f) from 86 sequences were used to midpoint point root the tree and resolved major tomato clades within the MYB-encoding genes (Fig. 6). The ML model and clustering analysis of *Aft* sequence grouped accessions into their expected clades with 60.4% bootstrap support for the separation of red-fruited species and green-fruited species (Fig. 6). The purple fruited *S. chilense* accession LA1996, clusters with other members of the green fruited clade and close to a purple fruited *S. habroachites* accession LA1777, with 44.7% bootstrap support (Fig. 6). Our purple accession *S. galapagense* accession LA1141 clusters with other members of endemic Galápagos tomatoes with 79.7% bootstrap support (Fig. 6). LA1141 does not cluster with members of the green-fruited clade based on sequence homology within the MYB-encoding genes underlying *Aft* (Fig. 6).

**Fig. 6.**
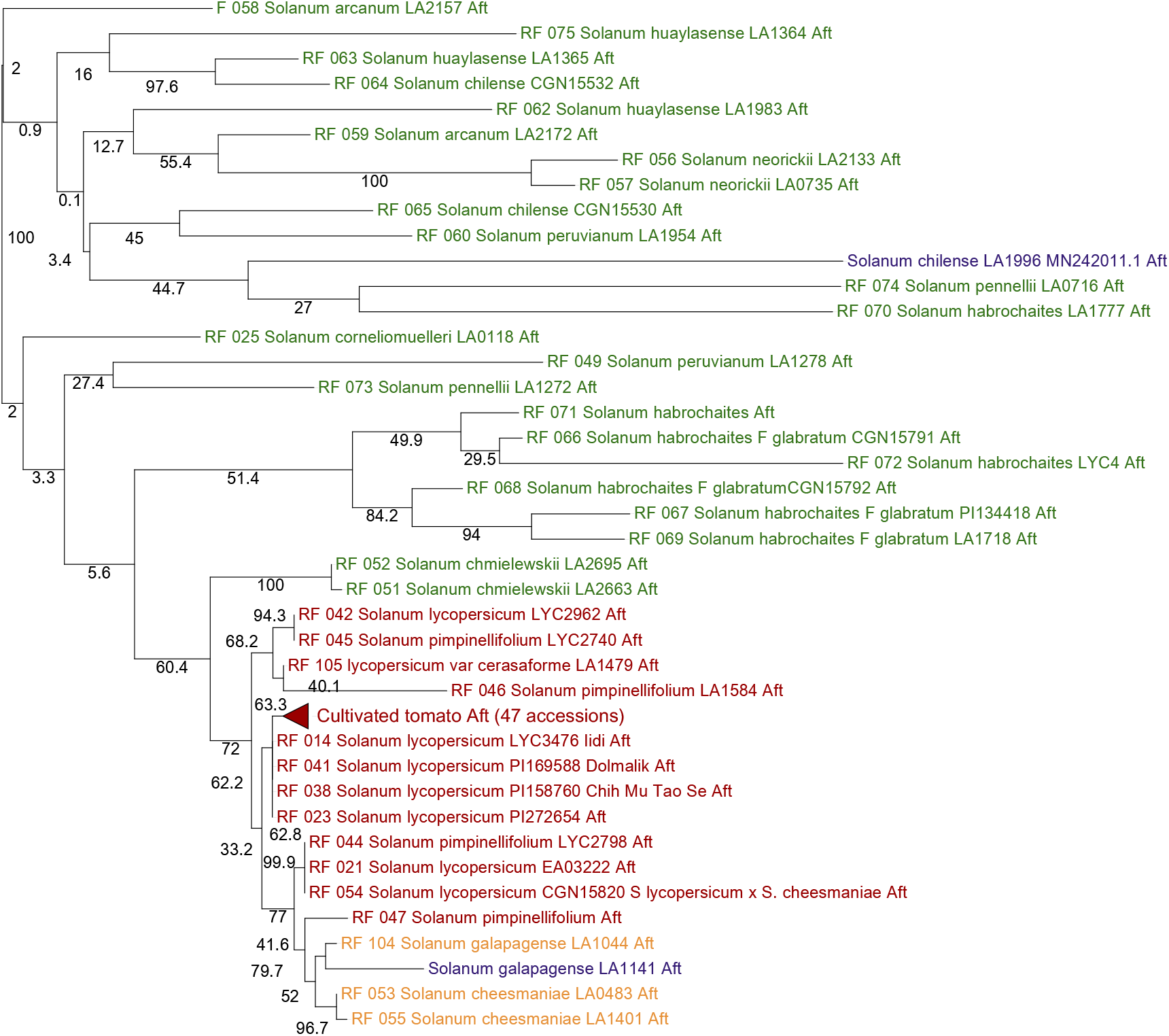
Midpoint rooted phylogenetic tree for MYB transcription factors underlying the *Aft* locus. The tree represents clustering of genomic sequences underlying *Aft* 84 unique tomato accessions from the 100 Tomato genome sequencing consortium, LA1996 (purple), OH8245 and LA1141 (purple) are clustered. A maximum likelihood midpoint rooted tree was constructed in the phangorn R package using the G.T.R model. Data resampling using 1000 rapid bootstrap replications was performed using the boostrap.pml function and bootstrap values are given for each branch. There are 47 identical *S. lycopersicum* sequences are condensed under the name “Cultivated tomato Aft (47 accessions) (**red triangle**).

Additionally, we clustered the coding sequences (CDS) corresponding to the *Ant1* and *An2-like* MYB genes underlying *Aft* from LA1141, OH8245, and 84 re-sequenced accessions with outgroup sequences from *Arabidopsis thaliana, Saliva miltorrhiza, S. tuberosom, S, lycopersicoides, C. annum, S. chilense* accession LA1996, S chilense accession LA1930, and *S. lycopersicum* variety Indigo Rose (Fenstemaker et al., 2021g). The CDS corresponding to *Arabidopsis thaliana* MYB genes that were determined to be homologous to *Solanum Aft* sequence were used as an outgroup to root the tree (Fig. 7). The ML phylogeny separated *Ant1* and *An2-like* CDS with 98.4% bootstrap support (Fig. 7). *Arabidopsis thaliana* and *Salvia miltiorrhiza* clustered closer together compared to accessions of *Solanum* for both *Ant1* and *An2-like* (Fig. 7). These results are consistent with previously published asterid phylogeny (Zhang et al., 2020). Accessions of *C. annum* clustered further from *S. tuberosom* (Fig. 7), consistent with *Solanum* phylogeny (Särkinen et al., 2013). For *Ant1* CDS, accession LA1141 clustered with members of the red fruited clade with 81.3% bootstrap support. For *An2-like* CDS, accession LA1141 clustered with members of the red-fruited clade with 49.7% bootstrap support (Fig. 7).

**Fig. 7.**
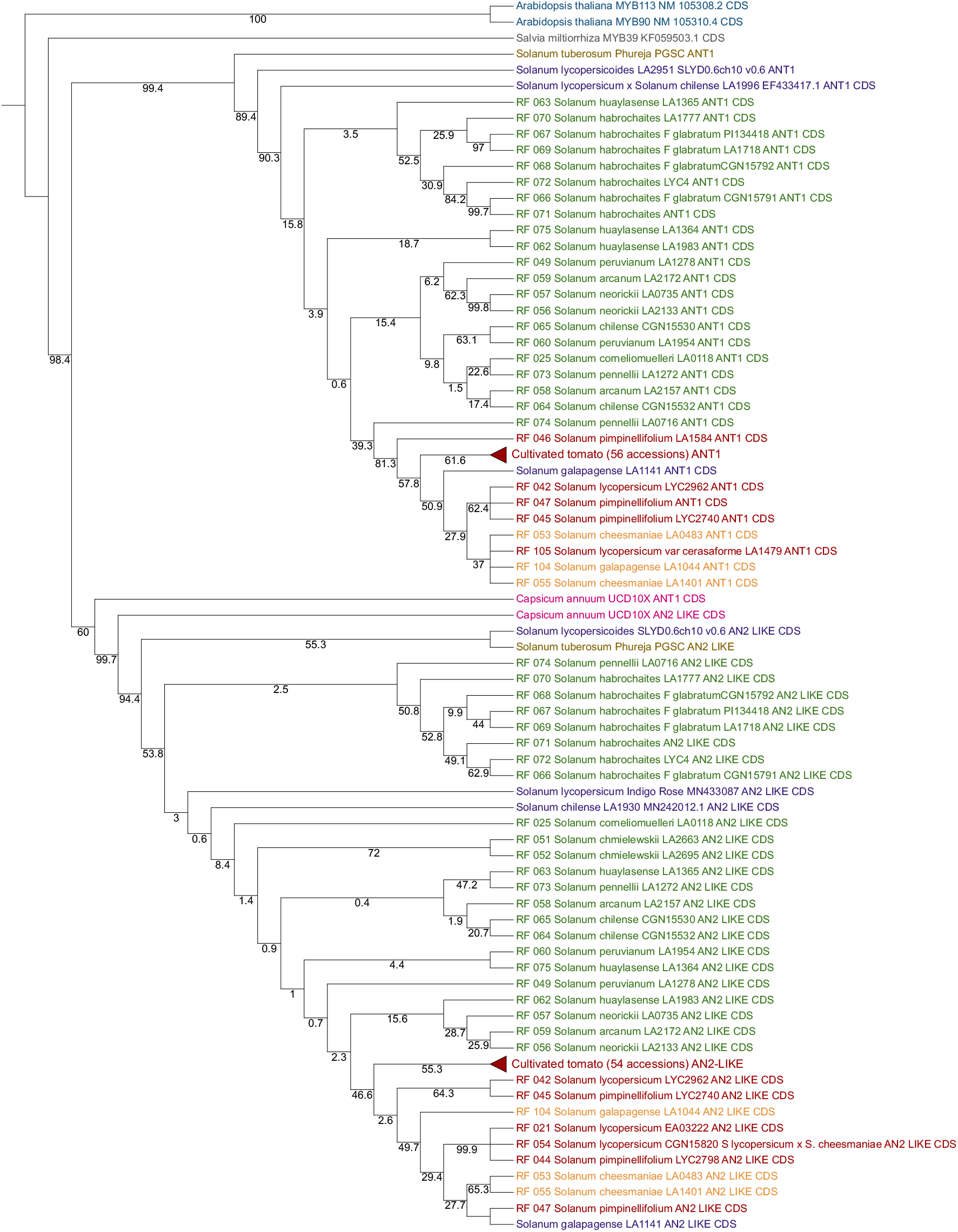
Outgroup rooted phylogenetic tree for MYB transcription factors underlying *Ant1* and *An2-like* coding sequence (CDS) at the *Aft* locus. *Arabidopsis thaliana, Salvia miltiorrhiza, S. tuberosome* Phureja, *C. annum, S. lycopersicum* ty Indigo Rose [MN433087 (Yan et al., 2020)], *S. chilense* accession LA1996 [MN242011.1, EF433417.1 (Sapir et 008; Colanero et al., 2020a)], *S. chilense* (Dunal) Reiche (formerly Lycopersicon chilense Dunal) accession LA1930 42012.1 (Colanero et al., 2020a)], 84 tomato accessions published as part of The 100 Tomato Genome Sequencing ortium (The 100 Tomato Genome Sequencing Consortium, et al., 2014), *S. lycopersicum* variety OH8245, and *S. agense* accession LA1141 are clustered. Identical *S. lycopersicum* sequences are condensed (**red triangles**). A mum likelihood tree was constructed in the phangorn R package (Schliep, 2011) using the G.T.R model. Data mpling using 1000 rapid bootstrap replications was performed using the boostrap.pml function and bootstrap s are given for each branch. Trees were rooted at Arabidopsis thaliana MYB-encoding genes as the outgroup.

## Discussion

### Measuring tomato fruit pigmentation with quantitative methods

Tomato color depends on the type and quantity of pigments synthesized in the fruits. Anthocyanins are responsible for the purple coloration of immature LA1141 fruit. Delphinidin-3-rutinoside and petunidin-3-(p-coumaroyl-rutinoside)-5-glucoside were the major anthocyanins identified. As fruit ripened, the predominant anthocyanin changed from petunidin 3-(coumaroyl)rutinoside-5-glucoside in the MG stage to malvidin 3(coumaroyl)rutinoside-5-glucoside in the breaker stage (Fig. 2). The chemical basis of pigmentation in progenies derived from LA1141 is consistent with those identified in introgression lines containing alleles from the green-fruited wild relatives (Jones et al., 2003). Phenotyping with quantitative measurements of color allowed us to distinguish classes of fruit that were useful for later genetic analysis. Cao et al., (2017) reported that it was difficult to distinguish marker-classes of *atv* with qualitative phenotyping, but we were able to detect differences in values of hue between homozygous and heterozygous genotypes (Fig. 4B). Additionally, our linkage analysis using quantitative measurements was able to distinguish classes and showed that *Aft* is necessary to recover light purple color in progenies (Fig. 1B). However, two unlinked loci are needed to recover the deep purple phenotype found in IBL selection SG18-124 (Fig. 1C). Inheritance of purple pigmentation in the progenies derived from LA1141 is consistent with patterns inherited from wild relatives in the green-fruited clade.

### Three putative QTL affect LA1141 fruit color

Color was associated with QTLs on chromosomes 7 and 10, and candidate genes were identified. The MYB-encoding gene family underlying the *Aft* locus maps to the distal arm of chromosome 10 and was associated with higher hue values. Two QTLs, one on the proximal arm and one on the distal arm of chromosome 10, were associated with chroma. The *Golden 2-like* transcription factor underlying the *uniform ripening* (*u*) locus maps to the proximal arm and mediated the brightness or dullness of the color. Accession LA1141 has a functional *Golden 2-like* allele underlying the *u* locus. The *u* locus is responsible for increasing chromoplast number, chlorophyll accumulation, and changing chromoplast distribution (Powell et al., 2012). This chlorophyll accumulation causes immature fruit to have patches of darker green color, especially where fruit are attached to the pedicel (Fig. 1D). Sequence analysis of MYB-encoding genes underlying the *Aft* locus suggested that LA1141 may have a functional R2R3 MYB activator which could explain its purple pigmentation in early stages of fruit development, as measured by hue, chroma, and L*. An allele of *atv* on chromosome 7 was detected based on interactions with *Aft* that increased pigmentation measured as hue (Fig. 4A). The QTLs and the interaction between chromosomes 7 and 10 were also validated in the subsequent IBL derived F_2_ generations (Fig. 4B).

Two QTLs were associated with L*, one on chromosome 6 and one on the proximal arm of chromosome 10. Only the QTL on chromosome 10 was validated in subsequent generations (Table 3). The region on chromosome 10 mapped to *u*. The *u* locus is likely affecting measurements of fruit darkness for similar reasons mentioned above. We expected the QTL on chromosome 6 to be associated with the *Beta* (*B*) locus. However, mapping *B* failed to support this locus as a candidate (Table 2). We were unable to identify a candidate for the QTL on chromosome 6 corresponding to L* in the IBC population. However, the QTL on chromosome 6 only explained 10% of the phenotypic variance compared to 35% of the variance explained by *u* (Table 2). Additionally, when we mapped *B* in the subsequent F_2_ populations we could detect association in only 1 of the populations (Table 3). In the SG18-200 derived F_2_ population, *B* was associated with hue, but not with chroma or L* (Table 3). We believe that our ability to detect *B* in this population is attributed to the monomorphic alleles for *atv* and *u* reducing the range of hue (Table 3).

### The primary regulatory mechanism for anthocyanin accumulation is conserved in LA1141

The interaction between chromosome 7 (*atv*) and chromosome 10 (*Aft*) in the LA1141 × OH8245 IBC population results in deep purple fruit (Fig. 1C). This interaction suggests that the role of synergistic MYB regulatory genes underlying loci on 7 and 10 is conserved between LA1141 and the green-fruited species. A complex of interacting MYB transcription factors, basic helix-loop-helix transcription factors (bHLH), and WD40 repeat domains (WDR), known as the MYB-bHLH-WDR (MBW) modulates anthocyanin accumulation in plants (Colanero et al., 2020b). The R2R3 MYB activators compete with the R3 MYB repressors for interaction with the bHLH transcription factor in the MBW complex (Colanero et al., 2020b). A CRISPR/Cas9 mediated silencing of MYB genes underlying the *Aft* locus suggested that only *An2-like* is needed for purple pigmentation in the peel of the tomato variety Indigo Rose (Yan et al., 2020). The same study showed that restoring function of *atv* in Indigo Rose reverts the coloration back to the light purple phenotype that we observed in SG18-200 (Fig. 1B) (Yan et al., 2020). Additionally, *atv* sequence targeted using CRISPR in the coding region of the second exon, where we observed the G to A SNP in LA1141, resulted in a loss of function of the R3/bHLH binding domain in LA1996 (Yan et al., 2020). This targeted mutation caused a purple phenotype that was similar to what we observed in our deep purple accession (Fig. 1C).

### Aft in LA1141 is likely a gain of function resulting from convergent or parallel mechanism

Pigmentation in the tomato clade of Solanum is considered a phylogenetic signal with the expression of carotenoids and anthocyanins separating the green fruited and red fruited clades (Gonzali and Perata, 2021). It is interesting to speculate about how LA1141 acquired its purple fruit pigmentation and how selection forces might maintain this pigmentation. One plausible explanation for selection and maintenance of pigmentation may be related to the role of fruit pigmentation in enticing organisms that disperse seed (Grotewold, 2006). AS an example, orange fruit are postulated to have a selective advantage on the Galápagos Islands as the result seed disperser color preferences (Gibson et al., 2021). An Investigation of known seed disperser preferences on the Galápagos islands and LA1141 fruit could elucidate a possible evolutionary mechanism, but more exploration is required. The duplication of MYB transcription factors in flowering plants in general and the locus of linked family members on chromosome 10 specifically provides opportunities for selection (Pickersgill, 2018).

In the red-fruited clade the structure of *Aft* phylogeny places *S. galapagense* accessions closer to *S. pimpinellifolium* and other red cultivated tomatoes, which is consistent with previously published *Solanum* phylogeny (Grandillo et al., 2011). We can separate the members of the red-fruited clade in the *Lycopersicon* group from *Arcanum, Eriopersicon*, and *Neolycopersion* groups in the green-fruited clade, but our phylogeny lacks the resolution to separate the green-fruited species within those groups (Fig. 6; Fig. 7). These results are also consistent with other studies (Peralta et al., 2008, The 100 Tomato Genome Sequencing Consortium). Additionally, results from the outgroup rooted tree using CDS from distantly related species suggests that the green-fruited clade is ancestral. Anthocyanin-mediated purple fruit appears to have been lost in the red-fruited clade. The gain of function at *Aft* in LA1141 has its origin in the red-fruited clade and is not likely an ancient introgression from a green-fruited progenitor.

## Conclusion

We identified an accession of *S. galapagense* that has purple pigmentation in the fruit. Anthocyanins are responsible for this color. Genes underlying the *atv, Aft*, and *u* loci are implicated as candidates for major QTL. The loci *atv* and *Aft* interact suggesting the same mechanism producing anthocyanins in the green-fruited clade is responsible for pigment patterns in LA1141 fruit. *Aft* is known from wild accessions in the green-fruited clade and we probed Rick’s hypothesis about an ancient hybridization event between progenitors of *S. galapagense* using genomic sequence from the *Aft locus*. Our phylogenetic analysis concluded that a functional allele of *Aft* in LA1141 is likely the result of convergent or parallel mechanisms and is not derived from introgression from a green-fruited relative. Our findings guide us toward a better understanding purple color found in the endemic Galápagos tomatoes and provide additional resources for characterizing anthocyanin biosynthesis in wild tomato relatives.

## Acknowledgements

We thank Jihuen Cho and the farm crews from the Ohio Agricultural Research and Development Center (OARDC) Wooster for assistance with management of the research. We thank Marcela Carvalho Andrade, Regis de Castro Carvalho, and Wilson Roberto Maluf from The Federal University of Lavras, 37200-000 Lavras, Brazil for assistance with the LA1141 IBC population. Salaries and research support were provided by state and federal funds appropriated to The Ohio State University, OARDC, Hatch project OHO01405, and grant funds from USDA Specialty Crops Research Initiative Award number 2016-51181-25404.

## Author contributions

SF and DF: conceptualization, SF: phenotyping, JC: chemical analyses, SF: linkage map construction, SF: QTL mapping, SF and LS: marker development and sequencing, SF: bioinformatics and sequence analysis, SF: phylogenetic analysis, SF: writing, and DF: contribution to writing

## Data availability

All data supporting the findings of this study are available within the paper. Additionally, pertinent supplementary tables and FASTA files are available in Zenodo at:

